# CheSPI: Chemical shift Secondary structure Population Inference

**DOI:** 10.1101/2021.02.20.432095

**Authors:** Jakob Toudahl Nielsen, Frans A.A. Mulder

## Abstract

NMR chemical shifts (CSs) are delicate reporters of local protein structure, and recent advances in random coil CS (RCCS) prediction and interpretation now offer the compelling prospect of inferring small populations of structure from small deviations from RCCSs. Here, we present CheSPI, a simple and efficient method that provides unbiased and sensitive aggregate measures of local structure and disorder. It is demonstrated that CheSPI can predict even very small amounts of residual structure and robustly delineate subtle differences into four structural classes for intrinsically disordered proteins. For structured regions and proteins, CheSPI can assign up to eight structural classes, which coincide with the well-known DSSP classification. The program is freely available, and can either be invoked from URL www.protein-nmr.org as a web implementation, or run locally from command line as a python program. CheSPI generates comprehensive numeric and graphical output for intuitive annotation and visualization of protein structures. A number of examples are provided.

## Introduction

NMR chemical shifts are very sensitive to the local structure of proteins, and can be measured and assigned routinely with great precision for both structured and unstructured proteins(Felli and Pierattelli 2012, Brutscher, Felli et al. 2015). The relationship between chemical shifts and local protein structure is well-established for folded proteins e.g. through simple index methods(Wishart, Sykes et al. 1992, Wishart and Sykes 1994) statistic and probabilistic methods(Eghbalnia, Wang et al. 2005, Wang, Chen et al. 2007) or by methods relying more on sequence homology through neural networks or database searches(Jones 1999, Labudde, Leitner et al. 2003, Shen and Bax 2013, Shen and Bax 2015), and super-secondary structure predictions(Hafsa, Arndt et al. 2015). Furthermore, CSs have also been used to aid the structural and dynamical characterization of proteins(Wishart and Sykes 1994, Wishart and Case 2001, Wishart and Case 2002, Berjanskii and Wishart 2007, Cavalli, Salvatella et al. 2007, Mielke and Krishnan 2009, Kjaergaard and Poulsen 2012, Robustelli, Stafford et al. 2012).

In stark contrast, intrinsically disordered proteins and regions (IDPs and IDRs) display no or very little regular secondary structure, are not folded into a globular structure, but rather constitute a dynamical equilibrium between several conformations with less regularity. NMR spectroscopy is an ideal technique to study these dynamic IDPs(Tompa and Fersht 2009, Uversky and Longhi 2010). Intrinsically disordered polypeptides display population-averaged chemical shifts that provide an operational definition of random coil chemical shifts. Deviations from RCCSs contain information about structural composition, but also pose a challenge to deconvolute induced from intrinsic structure. Fortunately, however, RCCSs have been predicted with increasing accuracy over time(Braun, Wider et al. 1994, Wishart, Bigam et al. 1995, Schwarzinger, Kroon et al. 2001, De Simone, Cavalli et al. 2009, Tamiola, Acar et al. 2010, Kjaergaard, Brander et al. 2011), culminating in the most accurate predictor to date, known as POTENCI(Nielsen and Mulder 2018). As a result, deviations from RCCSs (the secondary chemical shifts, SCSs) are now common parameters used to identify and quantitate order/disorder in IDPs(Berjanskii and Wishart 2007, Kjaergaard and Poulsen 2012, Nielsen and Mulder 2016, Sormanni, Piovesan et al. 2017). The improved accuracy has also permitted a benchmark of the performance of disorder prediction methods(Nielsen and Mulder 2019) and CSs were recently used to train the disorder predictor ODINPred (Dass, Mulder et al. 2020).

A yet more challenging task is to quantify the statistical composition of structural states for IDPs, since accurate reference experimental data with *unique* structural interpretation do not exist; As this problem is *ill-posed*, astronomical numbers of ensembles could be constructed that all give rise to the experimentally observed averages. One possible avenue to plausible solutions is to use physics-based models of protein conformational sampling (by e.g. molecular mechanics force fields) coupled to parametrization of chemical shifts, and possibly other NMR observables, to conformation(Ozenne, Schneider et al. 2012, Varadi, Vranken et al. 2015). Alternative empirical approaches adopt a heuristic treatment of CSs and secondary structure relationships in folded proteins to IDPs for the inference of secondary structure populations of α-helix, β-strand, random coil, and polyproline II(Camilloni, De Simone et al. 2012). A more robust reductionist approach was taken with SSP (secondary structure propensity)(Marsh, Singh et al. 2006, Tamiola, Acar et al. 2010) where linear combinations of SCSs were aggregated into a scale between −1 (sheet) and 1 (helix) and interpreted as a structural propensity. Although such an approach implicitly remedies correlated CSs, information potentially contained in the individual CSs risks being lost by the reduction to single value. For example, a rigid loop between a sheet and a helix will display near-zero propensity, and risks being falsely interpreted as disordered. Furthermore, neither of the above methods can so-far discriminate between disordered and ordered turns, whereas such information is potentially contained in the particular combination of SCSs.

Here, we introduce the linear analysis of signed SCSs, introducing **Che**mical shift **S**econdary structure **P**opulation **I**nference (CheSPI). This approach extends the previously-introduced CheZOD Z-score for quantifying local order and disorder in proteins(Nielsen and Mulder 2016, Nielsen and Mulder 2020) derived from the statistical analysis of sums of squared SCSs. Technically, CheSPI applies multivariate analysis and dimension reduction techniques to generate linear combinations (CheSPI components) of SCSs that optimally describing the variance: The first CheSPI component ensures the optimal distinction between secondary structure classes, whereas the second component accounts for the variance within the classes (which was found to be closely related to the local structure such as backbone conformation, flexibility and hydrogen bonding). CheSPI components offer an accurate and comprehensive quantification of local dynamic and structural composition in structured as well as disordered proteins, being sensitive to local protein structure as well as dynamics. CheSPI components are presented to the user as a color scale, which conveys the information in a simple, intuitive, and visually appealing manner. As shown herein, CheSPI colors can be used to annotate 3D structures, and thereby highlight important and detailed structural changes in proteins.

The power of CheSPI to discriminate between secondary structure classes was exploited to derive estimates for the populations of helix, extended structure, turn, and “non-folded” structures (CheSPI populations) through statistical inference, and these populations were validated through comparison to simulated ensembles for four distinct IDPs (*vide infra*). Contemporary methods for inferring secondary structure from NMR-data typically provide only three-class predictions. A notable exception is CSI3.0, which offers a four-state prediction including turns, as well as the distinction between internal and external strands. CheSPI takes a stride further, and extends secondary-structure inference to encompass the prediction of the eight structural classes (SS8) defined by the popular DSSP classification algorithm(Kabsch and Sander 1983); α-helix, 3_10_-helix, π-helix, extended β-strand, bridge (isolated single residue β-strand), and – if no well-defined pattern can be found – “bend” and “coil”.

### Availability

CheSPI is available at www.protein-nmr.org and source code can be obtained from https://github.com/protein-nmr. The CheSPI analysis is summarized in text files as well as in a combined plot containing three panels visualizing for each residue along the sequence: (i) CheSPI colors and CheZOD Z-scores, (ii) CheSPI populations, and (iii) CheSPI 8-state secondary structure predictions.

## Results

### Derivation of CheSPI components for secondary structure

NMR chemical shifts (CSs) are very sensitive to the local structure and dynamics in peptides. To analyze this correspondence, we used two previously derived sequence databases with CSs for proteins deposited in the BMRB database. The first database contains primarily disordered residues used to parametrize POTENCI(Nielsen and Mulder 2018), whereas the other, derived from the RefDB database(Neal, Nip et al. 2003), contains primarily structured residues. Secondary chemical shift (SCSs) were derived by subtracting random coil shifts derived by POTENCI corresponding to the backbone atoms and Cβ, and Hβ For the disordered database, residue data were labeled with “D” for disorder if their CheZOD Z-score (derived by CheZOD(Nielsen and Mulder 2016)) was less than 5.0, or else “O” for order. Analysis with DSSP(Kabsch and Sander 1983) was performed for all proteins in the structured database, and here residue data were labeled using the 8-class DSSP secondary structure designations. Subsequently, a supervised modelling approach was applied for dimensionality reduction, which optimizes the discrimination between the different classes (or equivalently, maximizes their separation). This procedure, called Orthogonal Projections to Latent Structures Discriminant Analysis (OPLS-DA), resembles PCA analysis(Worley and Powers 2016), but quantifies the variance *between* the classes with the first principal component whereas the variation *within* the classes is captured in the other dimensions (https://www.sartorius.com/en/knowledge/science-snippets/explaining-differences-or-grouping-data-opls-da-vs-pca-data-analysis-507204). The optimal weights for the principal components were derived using Simca-P(Wu, Li et al. 2010) as defined in Eqs. 1–3, Methods.

Figure 1 shows the loading plot that visualizes the optimized weights, scaled according to gyromagnetic ratio for each nucleus. The relative magnitude of the optimized weights reflects the well-documented sensitivity of SCSs towards secondary structure (Hα/C’/Cα/Cβ ≫ N/Hβ/H_N_), with Hα the hydrogen most sensitive to structure, and the three ^13^C nuclei displaying a similar importance. In addition, N/H_N_/Hα/Cβ display opposite SCSs compared to C’/Cα Hβ SCSs correlate with secondary structure as commented in Supplementary Results 1 and Supplementary Figure 1. For the second component, all weights are positive and display the following sensitivity: (H_N_ > C’/Cα/Cβ > N > Hα > Hβ). It is noteworthy, that the two chemical shifts with the largest magnitudes, H_N_ and C’, were previously found to undergo the largest chemical shift changes upon backbone hydrogen bonding(Nielsen, Eghbalnia et al. 2012). The third CheSPI component did not add any significant value to classification.

**Figure 1:**
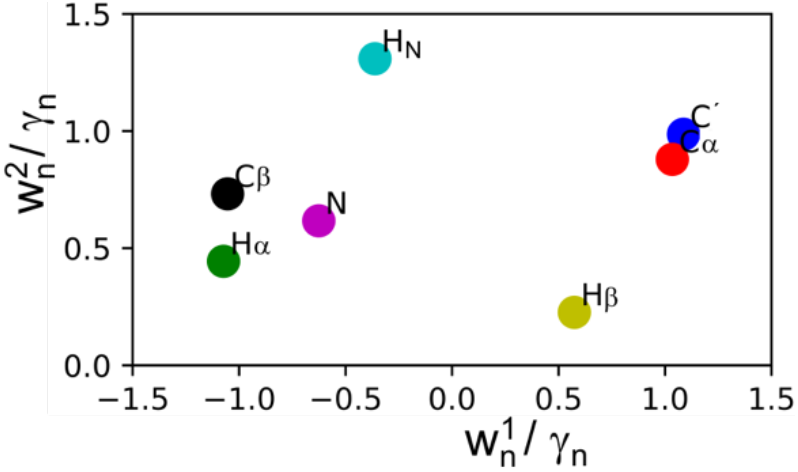
CheSPI component loading plot showing the weights (Eq. 1 Methods) scaled by gyromagnetic ratio for each nucleus.

### CheSPI components discriminate between local folded structures

CheSPI components were calculated for the 809 proteins in the structured database using the weights optimized by the OPLS-DA procedure (Eqs. 1–3, Methods). Figure 2 shows two-dimensional histograms of the combined observation of the first two CheSPI components for the three canonical secondary structure types helix, sheet, and coil as determined by DSSP. It is clear how the first component (*P*^1^) offers a near-perfect discrimination between helix and sheet whereas coil, placed around the middle of the score plot, overlaps with the helix and sheet classes, but has a lower average value for the second CheSPI component compared to helix/sheet (see also Supplementary Results 1 and Figure S1). For reference, it is noted that disordered residues have near-zero values for both of the first CheSPI components, as expected

**Figure 2:**
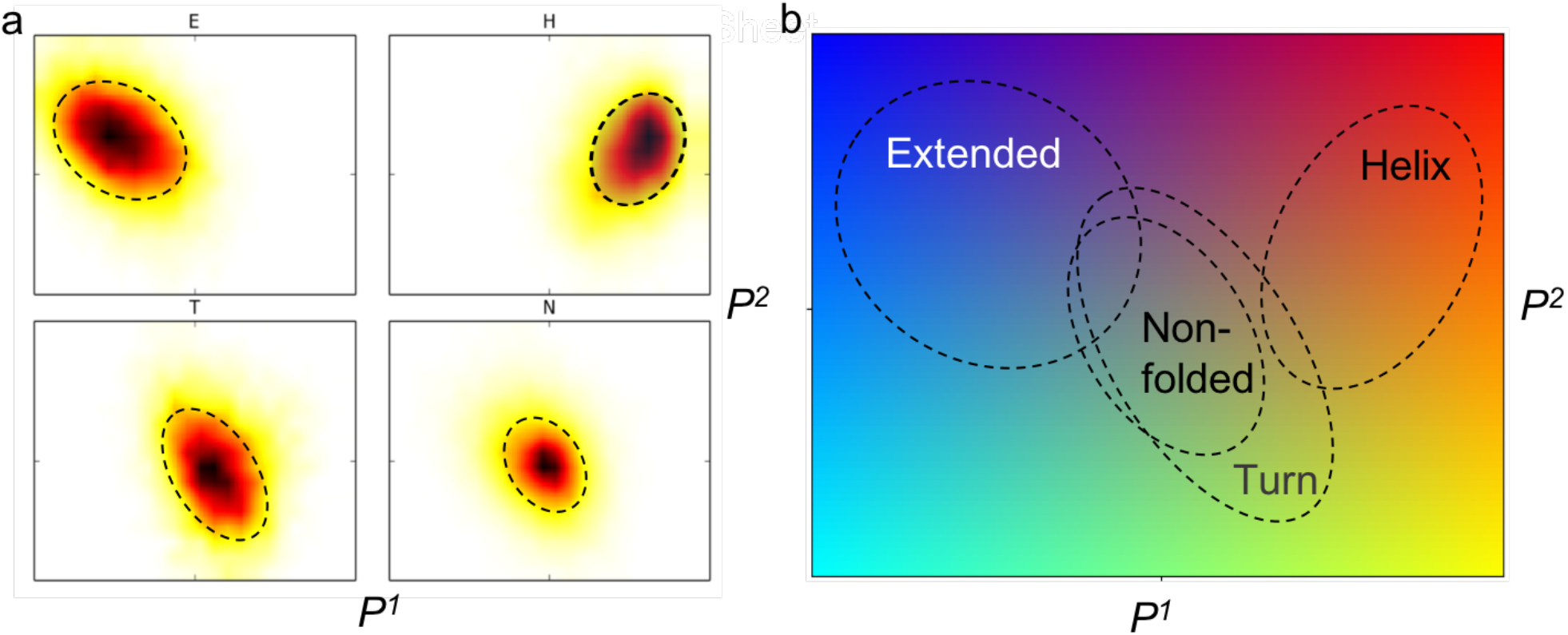
(**a**) Experimental distributions of the first two CheSPI components (Eq. 1–3, Methods) for the secondary structure classes; helix (H), extended (E), and turn (T) and non-folded backbone conformations (N, everything not helix, sheet, or turn) with ellipsoids marked by dashed lines. (**b**) overlay of ellipsoids from (a) merged with CheSPI color scheme.

Encouraged by the strong relationship to local structure, CheSPI components were converted to a color scale, using linear combinations of the first two components (see Eqs. 4 and 5, Methods). This CheSPI color scale provides a visual interpretation of the CheSPI components and an intuitive overview of the local structure and dynamics of proteins. The CheSPI application produces both bar plots and a script for 3D structure visualization based on CheSPI colors and several examples are discussed below (see Discussion). On this scale, well-formed strands and helices are defined by blue and red colors, respectively, coil can display multiple colors depending on context, turns are in green, and disordered residues are grey. The variation in CheSPI components along the secondary elements, as described above, is reflected in hues changing from red through orange to green at the C-terminal ends of helices, and at the ends of β-strands, which sometimes have lighter blue or purple CheSPI colors (see Figure 4a for an example of CheSPI colors).

**Figure 3:**
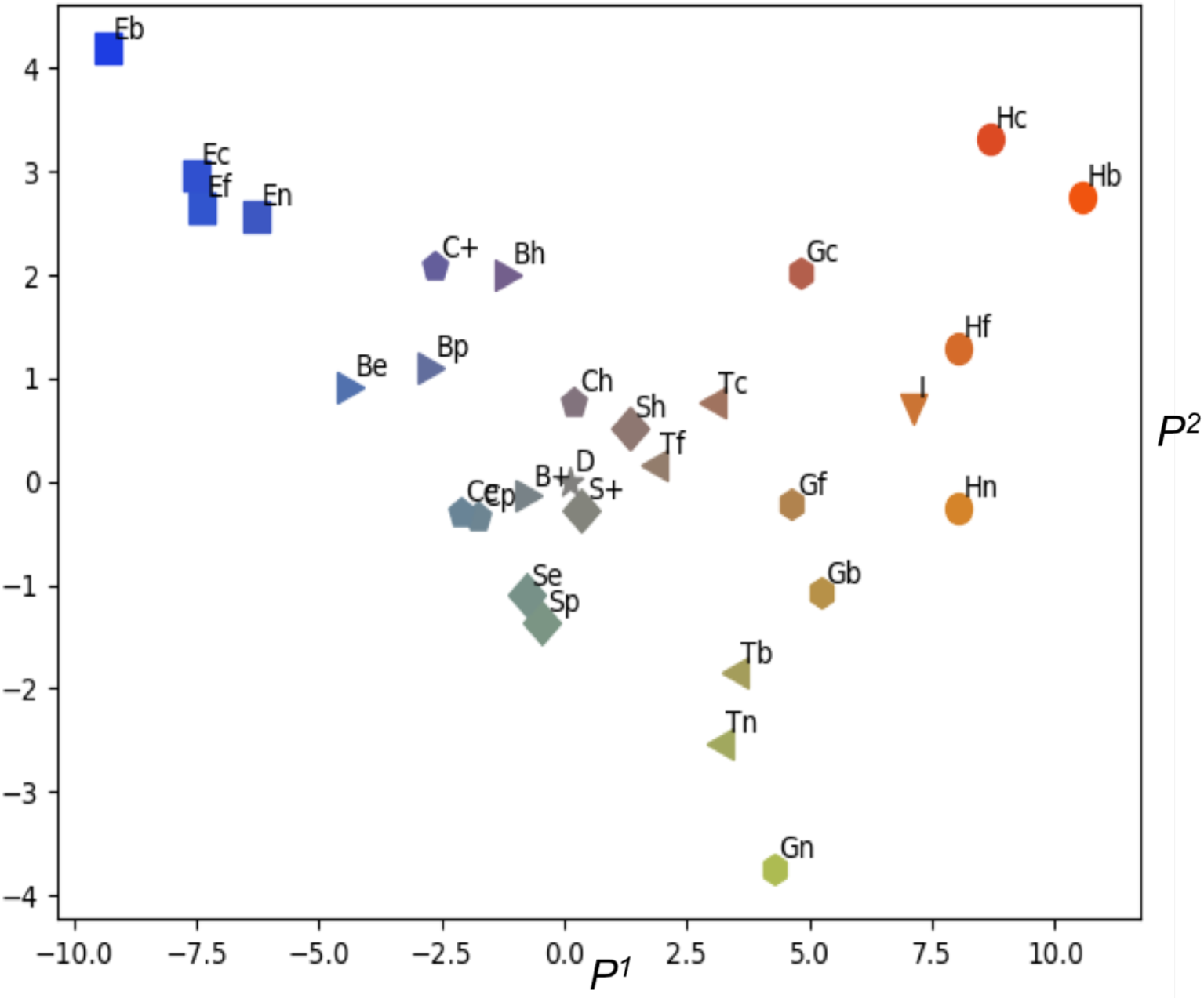
Average values of the first two CheSPI components for all 8 DSSP secondary structure classes with subdivisions. Average values were derived from the set of structured proteins. The average points are labelled with two letters (except I-helix and disorder), where the first letter indicates the DSSP classification (“C” indicates coil). The major classes E(β-sheet)/H(α-helix)/G(3_10_-helix)/T(turn) are labeled to indicate presence of hydrogen bonds (HB) using: n/c/b/f corresponding NH hydrogen bonding, C’-HB, both NH and C’, and none (free), respectively. For the remaining classes, B(bridge), S(bend) and C(coil), points are label according to the local backbone structure with: +/h/p/e corresponding to “positive”/”helical”/”poly-proline II”/”extended”, respectively, using the definitions in Table 1. The eight classes are shown with different marker symbols: box, circle, hexagon, triangle pointing left (left triangle), right triangle, down triangle, diamond, and pentagon, respectively for classes H/E/G/T/B/I/S/C and visualized using CheSPI color fills corresponding CheSPI components at the point (see Eq. 4 and 5, Methods). The corresponding point for disorder is indicated at the origin with a star, for reference.

**Figure 4:**
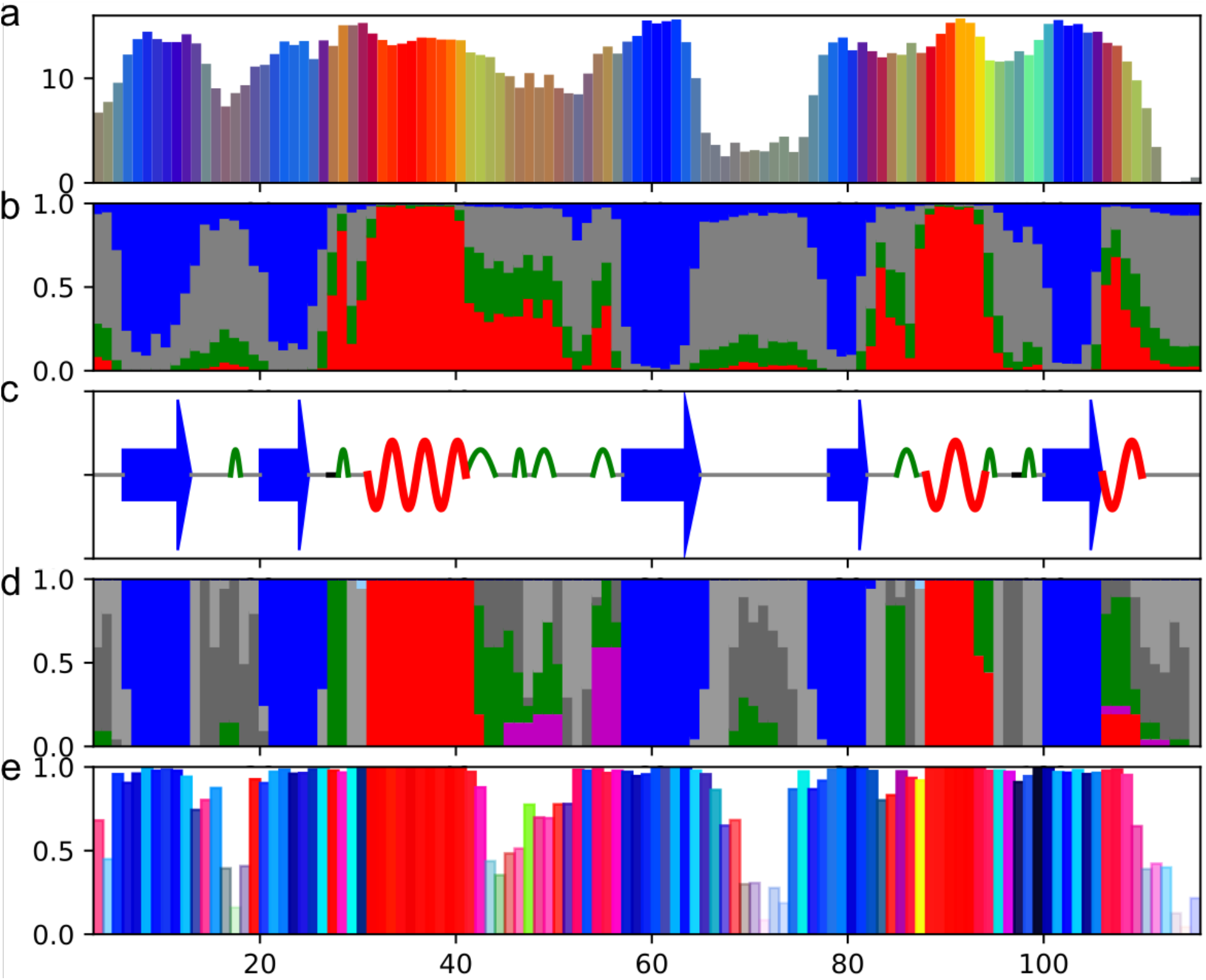
Output multi-panel plot produced by CheSPI using chemical shifts from BMRB id 6635 and validation from structural ensemble for PLCɛ-RA2(Hyberts, Goldberg et al. 1992) (PDB id 2BYF). (**a**) Bar plot color with CheSPI colors (Eq. 5 Methods) with bar height equal to the CheZOD Z-scores. (b) Stacked bar plot of CheSPI populations of “extended” (blue), “helical” (red), “Turn” (green), and “non-folded” (grey), local structures, corresponding to DSSP classes of sheet and bridge (E/B), all helix-types (G/H/I), turn (T) and, finally, the remaining bend and coil (S/-), respectively (see text). (c) Cartoon of the most confident CheSPI prediction of eight class DSSP secondary structure using red curved lines for α-helix (H), magenta for 3_10_-helix (G), blue arrows for sheets (E) and bridges (B), green arcs for turns (T), and grey and black lines for coil (C) and bend (S), respectively. (d) Stacked bar plot visualizing the observed conformations in the structural ensemble for the 8 DSSP classes using same colors for helixes, turns, and lighter and darker grey colors for coil and bend, respectively. (e) Bar plot for backbone angle conformation and variation. The heights of the bars were set to the geometrical average of the squared angular order parameters(Hyberts, Goldberg et al. 1992) for ϕ and ψ backbone torsion angles (Eqs. 14–16, Methods) where values close to unity indicate local order and values close to zero indicate structural disorder (large angular variation). The colors were taken from a 2D-color-scale (see Figure S3) based on the position in Ramachandran plot of pairs of ϕ and ψ backbone torsion angles using trigonometric averages of the ensemble values (see Eq 17, Methods). With this scale, backbone angles in the helical domain of the Ramachandran map appear in red as before, and extended β-sheet-like conformations have blue colors. Furthermore, left-twisted b-strands as well as fragments with PPII structure appears with cyan colors whereas conformations with positive ϕ have yellow and green colors, and finally, other conformation referred to elsewhere as “forbidden” in Ramachandran space are shown with black colors. Transparency is added to the bars using the above local angular order parameters as the “alpha value”. See also Figure S1.

To further demonstrate the power of the CheSPI components to discriminate between different local structures and describe variation within structures, we analyzed the eight DSSP classes. These were further separated into subclasses based on hydrogen bonding or local backbone structure as visualized in Figure 3 (see also Fig. S2 for individual histograms). Average values of the first two CheSPI components for all subclasses are visualized in Figure 3. First, it is seen how the first CheSPI component (*P^1^*) roughly segregates the average values for the 8 DSSP classes with E<B<C<S<T<G<I<H, with the more extended conformations having the most negative values, in general. Secondly, the second CheSPI component (*P^2^*) shows negative values for turns and positive ones for helices and strands. Notably, *P^2^* accounts for the variation within the same DSSP class. For the helix and turn classes, it appears that in case of H_N_, hydrogen bonding reduces the value of *P^2^*, whereas for C’, H-bonding increases it. This tendency reflects the larger contribution from H_N_ and C’ chemical shifts in the definition of *P^2^* as described above. Conversely, the effect of hydrogen bonding is less pronounced for *P^2^* in the case of strands. Here, strands with both H_N_ and C’ hydrogen bonding (“Eb”) have larger magnitudes for both CheSPI components. Such “Eb” strands correspond to the inner strands of β-sheets and are identified by deeper blue CheSPI color (see Discussion below). The variation in *P^2^* within strands likely relates to their variability in local structure, including twists, bends, and bulges, as witnessed by examples given below. Finally, the classes with lower tendencies to form hydrogen bonds, B(bridge), S(bend) and C(coil), have CheSPI components closer to zero. The more extended conformations “extended” and “poly-proline II” (PPII) (see legend to Figure 3) show lower values for *P^1^*, while the more compact conformations, “helical” and “positive ψ”, display larger values for *P^2^*. Markedly, “extended” and PPII conformations have almost indistinguishable CheSPI components for bend and coil, whereas for bridges, “extended” displays slightly lower values for *P^2^* in agreement with the closer resemblance to canonical β-sheet.

### Prediction of secondary structure populations

Intrinsically disordered proteins are in a dynamic equilibrium between different local conformations. Encouraged by the discrimination power of the CheSPI components (Figs. 2 and 3), we derived an estimate for the population of different local structural states based on the relative likelihood for observing a given combination of the first two CheSPI components as defined in Eq. 6 in Methods. **Che**mical shift **S**econdary structure **P**opulations **I**nference (CheSPI) are provided for classes referred to here as “extended”, “helical”, “turn”, and “non-folded”. This classification is based on inference from statistics of CheSPI components measured in the structured proteins set for DSSP classes for strand and bridge (E/B), all helix-types (G/H/I), turn (T) and, finally, the remaining non-folded bend and coil (S/-), respectively.

**Table 1:**
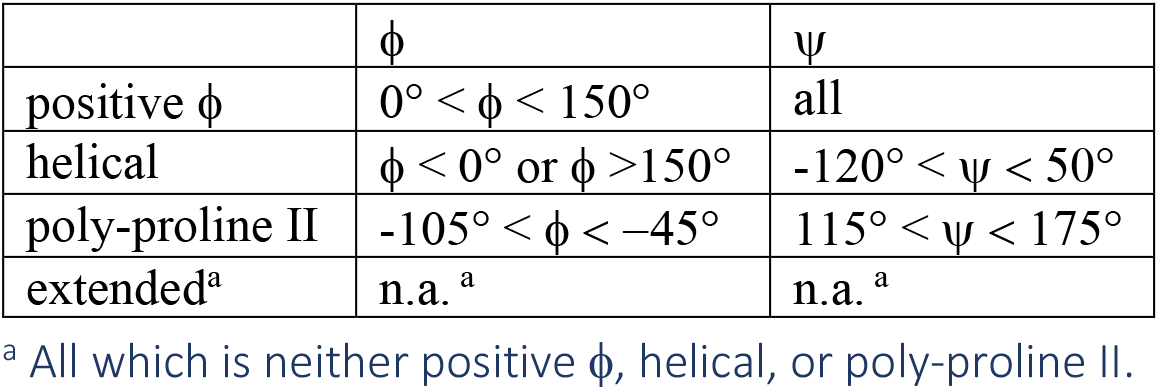
Definitions of local backbone conformations.

We now turn our attention to a small number of examples for demonstrating the utility of CheSPI. First, we analyze in detail the NMR solution structural ensemble of the phospholipase c epsilon RA 2 domain (Hyberts, Goldberg et al. 1992) (PLCɛ-RA2, henceforth) as summarized in Figure 4 (see also Figure S1). PLCɛ-RA2 contains 5 β-strands, two α-helices and two shorter 3_10_-helices which are modelled in some of the members of the deposited NMR ensemble (PDB id 2byf). The α-helices are predicted with close to 100% population throughout. Although for the β-strands, populations of about 90% “extended” were predicted for the central residues, relatively lower estimates are obtained at both ends of the strands, echoing the fall-off in CheSPI component amplitudes at β-strand ends described above. At the same time, significant populations for “extended” were predicted in loop segments for residues flanking the β-strands, mirroring the gradual change from strand character to flexible coil, bracketing β-sheets. More interestingly, disordered regions are characterized by a composition of different local structures, and the termini and the long loop region (residues 66-76) and classified as disordered, when judged by CheZOD Z-scores < 8.0. For these regions, larger “non-folded” populations (> 70%) were indeed predicted by CheSPI. Of note, residues 68-76 have missing density in the corresponding X-ray structure(Bunney, Harris et al. 2006) (pdb id: 2C5L) suggesting that they may be dynamically or statically disordered. In the corresponding NMR structure ensemble (Fig. 5) (Hyberts, Goldberg et al. 1992), a mixture of bend and coil conformations as well as a few turns are observed for this long loop. Averaging of the local backbone conformations in this loop leads to small angular order parameters(Hyberts, Goldberg et al. 1992, Nielsen and Mulder 2019) (Fig. 4e) indicative of increased local disorder. Average chemical shifts for the individual conformations lead to observations very close to random-coil chemical shifts for these residues, and thereby low CheZOD Z-scores. Interestingly, for the other long loop (residues 45-56), CheSPI estimated intermediate populations of α-helix (ca. 30-40% for residues 45-50 and 54-55), and comparison with the NMR ensemble reveals fractional populations of two 3_10_-helices for residues 45-50 (15-20% of the members in the ensemble) and residues 54-56 (60%).

**Figure 5:**
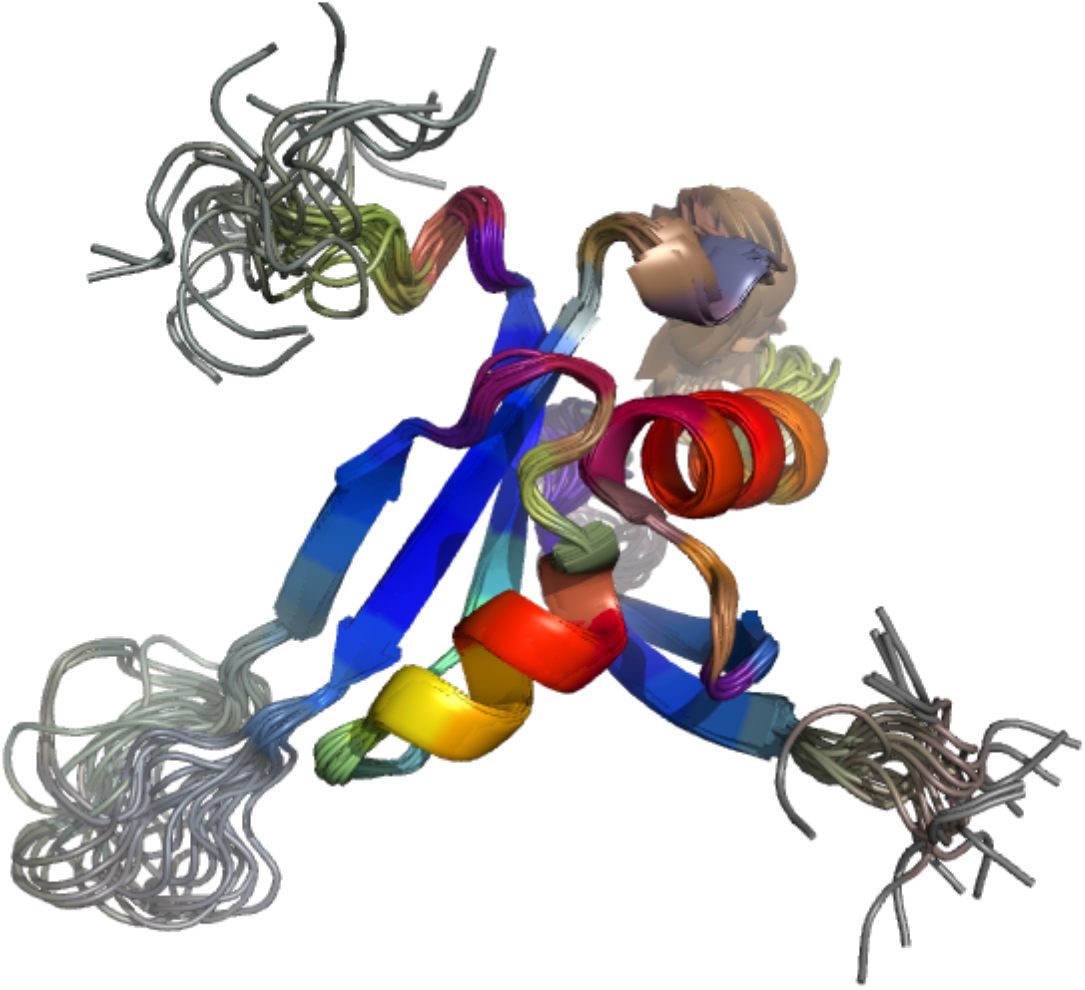
**Structure visualization using CheSPI colors** for the NMR solution structural ensemble of PLC025B;-RA) (Hyberts, Goldberg et al. 1992) using chemical shifts from BMRB id 6624 and PDB id, 2BYF.

### Validation of canonical secondary structure populations in disordered protein ensembles

Although the range of conformations in an ensemble of structures determined by NMR spectroscopy in solution are evidence of the dynamics of the system, the generated models also reflect the precision of the structure, and depend on the type, quality and number of geometrical constraints as well as the structure determination protocol. More advanced computational protocols(Bernadó, Blanchard et al. 2005, Jensen, Salmon et al. 2010, Marsh and Forman-Kay 2012, Ozenne, Schneider et al. 2012, Camilloni, Cavalli et al. 2013, Varadi, Vranken et al. 2015) use extensive conformational sampling, culled by data from NMR spectroscopy and small-angle X-ray scattering (SAXS). To gauge how well CheSPI-derived populations compare with the composition of local structure, we made comparisons for four protein systems: (i) The K18 domain from Tau, a human intrinsically unstructured protein implicated in Alzheimer’s disease pathology(Cleveland, Hwo et al. 1977). K18 was previously investigated by NMR chemical shifts, residual dipolar couplings (RDCs), and paramagnetic relaxation enhancements (PREs)(Mukrasch, Markwick et al. 2007), and an ensemble of structures was computed(Ozenne, Schneider et al. 2012) using a combination of the ASTEROIDS(Jensen, Salmon et al. 2010) and Flexible Meccano(Bernadó, Blanchard et al. 2005) protocols (see also Discussion); (ii) The unfolded state of drkN SH3. A structural ensemble of this small domain has been generated with the ENSEMBLE software(Marsh and Forman-Kay 2012) based on an extensive amount of experimental data(Marsh and Forman-Kay 2009) including CSs, RDCs, SAXS, PREs, and ^15^N R_2_ relaxation data. Here we compare CheSPI populations based on the assigned chemical shifts(Lee, Zhang et al. 2015) (iii) The PaaA2 antitoxin. This protein contains two partially formed helices, and was modelled based on a combination of NMR data, filtering with SAXS, and cross-validation with RDCs(Sterckx, Volkov et al. 2014). These three ensemble structures were taken from the pE-DB protein ensemble database for IDPs(Varadi, Kosol et al. 2014). (iv) The oncogene protein E7 of human papillomavirus type 16(Kukic, Lo Piccolo et al. 2019), which contains both ordered and disordered domains with an ensemble of conformations similulated by replica-averaged metadynamics (RAM) simulations(Camilloni, Cavalli et al. 2013) based on assigned CSs and RDCs. In all cases, the local secondary structure was calculated by DSSP for each member of the ensemble. The latter was stratified into populations of helix (H/G/I DSSP classes), β-strand (E/B), and a coil class. “Coil” was further divided according to the backbone conformations as described above (see legend to Table 1).

A good correlation between “observed” fractions of formed helix conformations in the ensembles is seen in Figure 6, with Peason correlation coefficient 0.915, and a similar degree of correlation for strand structures (R = 0.911). For comparison, the δ2D algorithm predicts populations of helix and strand with correlations to the observed fraction of populations in the ensembles of 0.85 and 0.78 for helix and strand, respectively. Furthermore, ncSPC yields comparable correlations of 0.87 and 0.79 for helix and strand (Tamiola and Mulder 2012), respectively, when interpreting positive secondary structure propensities (SSPs) as helix fraction and the absolute of the negative SSPs as strand fraction. The first CheSPI component is closely related to the ncSPC scale and gives correlations of 0.91 and 0.89 to helix and strand populations, respectively, which is close to the aggrement between CheSPI populations and observed fractions. When considering residues with local “helical” backbone structure as helix, and similarily, residues with local “extended” backbone structure as strand, the correlations between predicted and observed population decrease (between R = 0.63 and 0.77) but the ranking of the performance of the methods is preserved.

**Figure 6.**
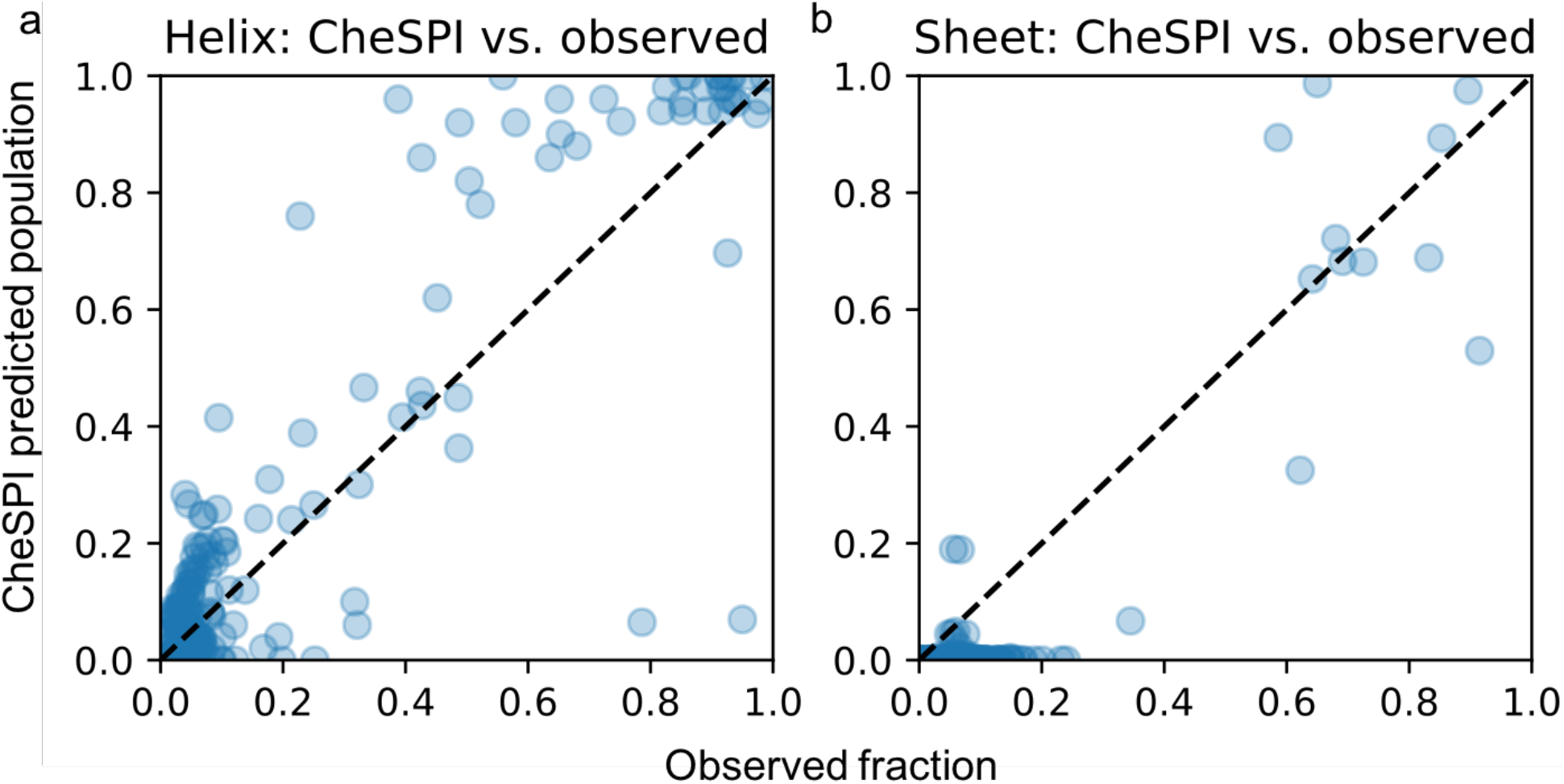
Correlation between helical populations predicted by CheSPI and observed in experimental ensembles. Each residue data is indicated with a blue disk. Populations were using data from (BMRB id, pE-DB id) = K18 Tau: (19253, 6AAC), PaaA2: (18841 5AAA), DrkN SH3 unfolded; (25501, 8AAC), E7: (BMRB id’s 19442 and 26069 for residues 3-45 and 46-97, respectively, using coordinate data provided by the authors).

### Prediction of secondary structure according to eight-class DSSP

CheSPI provides estimates for the populations of local structure types, whereas folded proteins are more commonly described by segments of completely formed regular structure. The classical DSSP algorithm assigns each residue to one of eight classes(Kabsch and Sander 1983), which are more informative about the local structure than the traditional coarse-grained three-state canonical division of secondary structure. Unfortunately, to date no tool exists that can infer this 8-class DSSP structure from NMR chemical shifts and sequence alone. Therefore, we extended the CheSPI analysis to the prediction of secondary structure segments and 8-class DSSP secondary structure (SS8) using only protein sequence and assigned NMR chemical shifts as input. As presented above, CheSPI reveals clear trends of secondary structure in its principal components. To quantify this dependence, we thus defined a linear approximation back-calculation of the CheSPI components (PCs) based on sums of contributions from the (four) nearest neighboring SS8 and amino acid types (see Eqs. 7–9 Methods), e.g. the first component would decrease if the center residue was strand and to smaller degree if the neighboring residues were strand. All weights were parametrized by linear regression, similar to the POTENCI implementation(Nielsen and Mulder 2018), using observed PCs derived from the structured database as targets and observed SS8 and sequence as input variables (see Methods). By this procedure, the first PC was predicted with an average error of 1.97, 3.63, and 2.94 for helix, strand, and coil, respectively (the full span of PCs is almost one order of magnitude larger). This prediction of PCs was used together with the average error to estimate a likelihood of observing any PC given an SS8 assignment and the sequence (Eq. 10 Methods). Bayes’ theorem was then used to “invert” the probabilities, i.e. to give the probability of an SS8 given the observed PCs. By this procedure, the prior probability for the secondary structure is “updated” by multiplying with the likelihood of observing the PCs given the secondary structure and sequence (Eq. 11 Methods). Finally, the predicted secondary structure is the combination of SS8 states for all residues that give the maximal posterior probability (Eq. 12, Methods). Unfortunately, SS8 states for a residue cannot be optimized univariately, since the value of predicted PCs depends on the nature of the neighboring SS8s. Therefore, the posterior probability maximization algorithm was implemented using a genetic algorithm (as in POTENCI) to produce populations of SS8 assignments for the full sequence. The SS8 predictions are rapidly calculated (2-3 seconds). Finally, the SS8 assignment from the population with the lowest energy is considered the best prediction, and the variation within the population at each site, along with the agreement between predicted and observed PCs, is used to estimate a confidence for each residue SS8 prediction.

For PLCɛ-RA2 (Figure 4), CheSPI detects all secondary structure elements and identifies the borders of these elements (Fig. 4c) with high accuracy when compared to the observed secondary structures (Fig. 4d) (all either exact locations, or one off, and a single case with two off). The disordered stretches are predicted as coil (majority). The fractionally-occupied 3_10_-helices are predicted as turns by CheSPI, which is not very surprising given the similarity between 3_10_-helices and type I β-turns(Shapovalov, Vucetic et al. 2019)

To derive a systematic evaluation of CheSPI SS8 predictions, a validation set was generated by considering all entries in the BRMB database published after the newest version of the RefDB database, which was used to derive our first structured database for optimizing the CheSPI weights and perform inference. We aimed for a small validation set, keeping only entries with (i) less than 30% sequence identity to all sequences in the structured database, (ii) all backbone chemical shifts available including Hβ, (iii) a high quality X-ray structure for the corresponding sequence (R < 2.0 Å) with exact sequence identity to the NMR study, (iv) no biasing conditions in either the derivation of the NMR assignments or the X-ray structure (e.g. standard buffer conditions as before(Nielsen and Mulder 2016) and no large ligands present). This procedure yielded 13 protein entries (see Table S1) with assigned chemical shifts, which were used to generate CheSPI SS8 predictions and comparison to the observed secondary structure classes, as calculated with DSSP from the high-quality X-ray structures. CheSPI achieves a good accuracy with 68.6% (between 53.2% and 80.6) correct 8-class predictions (Q8), and 84.6% correct for the classical three-class predictions (Q3) being an improvement by 2.7% relative to the 81.7% (Q3) for CSI 3.0 (see Table S1). Furthermore, considering only the CheSPI predictions with the highest confidence (28% of cases), CheSPI performs with 94% for SS8 accuracy. High confidence is typically found in the middle of the secondary structure elements and in long disordered loops, whereas lower confidence is more likely to be observed at the borders between regular secondary structure and loop elements.

## Discussion

In this paper, we have introduced CheSPI components – derived from NMR secondary chemical shifts – that provide an aggregate descriptor of local structure and dynamics for both structured and disordered proteins. CheSPI components estimate the populations of secondary structure, and are visualized using color, rather than the previously-published SSP and ncSPC procedures(Marsh, Singh et al. 2006, Tamiola, Acar et al. 2010, Tamiola and Mulder 2012), which present a scale bar to differentiate only the two most common secondary structure *propensities* (SSPs). The first CheSPI component is similar to the SSP scale in its power to discriminate between different secondary structures, and gives comparable values (see below). CheSPI, however, supersedes ncSPC by the introduction of a second component that affords to account for the variation *within* structural classes - and thereby offers a far more comprehensive discrimination of local structure and dynamics in proteins. Alternatively, secondary structure populations can also be predicted by δ2D in order to stratify residues as “helix”, “beta”, “poly-proline II” and “coil”. CheSPI takes this differentiation further by its sensitivity to discriminate distinct non-folded and folded “turn” coil types from NMR chemical shifts. This advance is possible, as CheZOD Z-scores facilitate the appropriate classification of non-canonical local structure and dynamics.

To feature the potential of CheSPI for detailed structural analysis using NMR chemical shifts, we provide a few examples below. These examples demonstrate that small, but important changes in solution structures can be characterized from NMR chemical shifts, which may otherwise be difficult or impossible to capture.

### Metal binding and aggregation of Cu/Zn superoxide dismutase 1

As a first example, we focus on CS data for the protein Cu/Zn superoxide dismutase 1 (SOD1)(Milani, Gagliardi et al. 2011) in the *apo* and metal-bound *holo* states. The misfolding of SOD1 is linked to familial amyotrophic lateral sclerosis (ALS)(Rosen, Siddique et al. 1993, Robberecht, Sapp et al. 1994). SOD1 contains a double β-sandwich structure, where two long loops that are disordered in the *apo* form become structured upon metal binding(Rakhit and Chakrabartty 2006, Teilum, Smith et al. 2009, Sirangelo and Iannuzzi 2017). In Figure 7, CheSPI analysis reveals a clear difference between the *apo* and *holo* forms. CheZOD Z-scores (Fig. 7a, b) confirm that the bound form is structured, whereas the two largest loops (IV and VIII) are unstructured in the *apo* form. The first CheSPI component (related to secondary structure propensity) remains close to zero and doesn’t change much between the two states of the protein. On the other hand, the second CheSPI component changes to negative for the bound state, indicative of the formation of turn structure. This change from non-folded to folded coil is apparent in the changes from grey to green on the CheSPI color scheme for loops IV and VIII (Fig. 7c,d). Structure determination(Banci, Bertini et al. 2002) clearly shows how metal-ion coordination induces folding of these two large loops (Figs 7h and 8), which become enriched in turn structure. Using NMR relaxation dispersion measurements, Teilum and co-workers identified a weakly populated exited state of apo-SOD1, which is believed to trigger deviant oligomerization(Teilum, Smith et al. 2009). They showed that the largest structural changes between the *apo* ground and exited states involves the flexible loops IV, VI, and VII, as well as β-strands 4, 5, 7, and 8. While native dimers form through association of pairs of β1, it was discussed how excited-state exposure of edge strands 5 and 8, which are protected by turn structures in the metal bound form, could initiate the oligomerization process. Extensive aggregation is avoided by negative design(Richardson and Richardson 2002) of β5 and β8, which are more twisted and less hydrogen-bonded (Fig. 8b,c), contain less β-sheet in the structural ensemble, which is slightly higher for the *apo* form (Fig. 7h), and have fewer canonical β-sheet backbone angles (Fig. 7i). This is reflected in the lower populations of extended structure for these strands estimated by CheSPI (panels e-g). In contrast, β1 is fully formed in the ensemble with canonical backbone angles (Fig. 7h,i). Non-native Inter-molecular contacts were identified for several residues in loop IV, in particular residues H63 and F64 (no data available for residues 65-66) through paramagnetic relaxation enhancement(Teilum, Smith et al. 2009). Intriguingly, CheSPI analysis pinpoints a small segment, residues 61-66, with noteworthy local order within this loop (Fig. 7a, c). Residues 61-63 show significant “extended” CheSPI populations, whereas residues 64-65 have elevated helical population (Fig. 7e). Whereas H63 forms a β-bridge in the metal-bound structure, flanked by residues with extended conformations, residues 65-66 mostly populate helical backbone conformations in the ensemble structure (Fig. 7h, i). Similarly, residues 132-137 form a helix within loop VII of the *holo* form, with a pronounced peak of helical populations that is mirrored in the *apo* structure (Fig. 7e). CheSPI analysis suggests that the locally ordered residues 65-66 might initiate non-native oligomerization through contacts within preformed extended structure.

**Figure 7:**
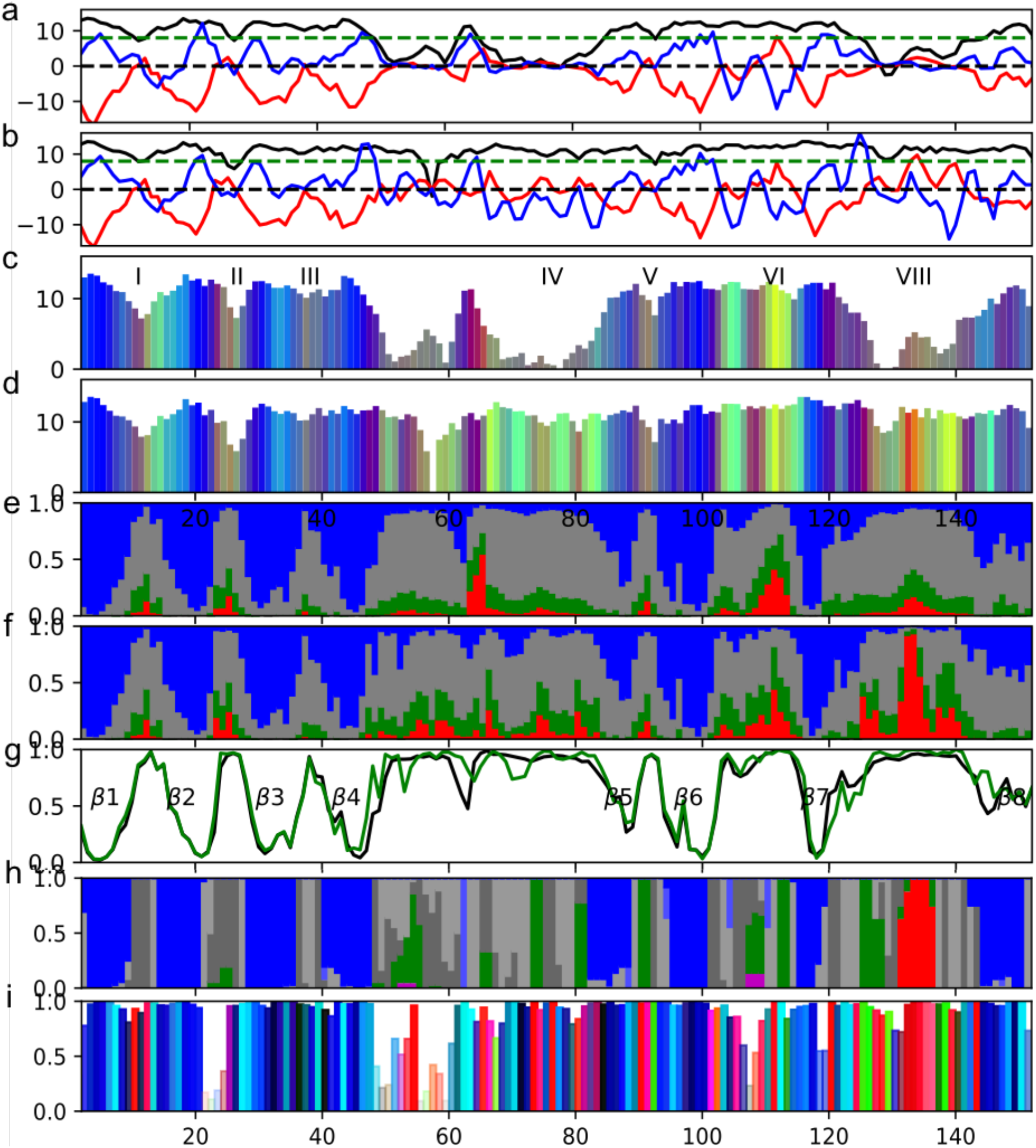
CheSPI analysis of Cu/Zn superoxide dismutase 1 (SOD1). (**a** and **b**) CheSPI components and CheZOD Z-scores for apo-SOD1 (see legend to Figure S1b) and using CSs from entry with BMRB id 15711 and Cu/Zn-bound state of SOD1 (BMRB 4402), respectively. (**c** and **d**) CheSPI colored bar plot for apo-SOD1 and Cu/Zn-bound SOD1, respectively (see legend to Figure 4a). Loops are labeled using roman numbers. (**e** and **f**) CheSPI populations for apo-SOD1 and Cu/Zn-bound SOD1, respectively (see legend to Figure 4b). (**g**) plot of CheSPI predictions for “extended” populations for apo-SOD1 (black curve) and Cu/Zn-bound SOD1 (green), respectively. β-strands are labeled consecutively with Greek letter and Arabic numbers. (**h**) Local structure observed population in structural ensemble for and Cu/Zn-bound SOD1 (PDB id 1ba9) (see legend to Figure 4d). (**i**) Average backbone angles and angular order parameters for Cu/Zn-bound SOD1 (PDB id 1ba9) (see legend to Figure 4e).

**Figure 8:**
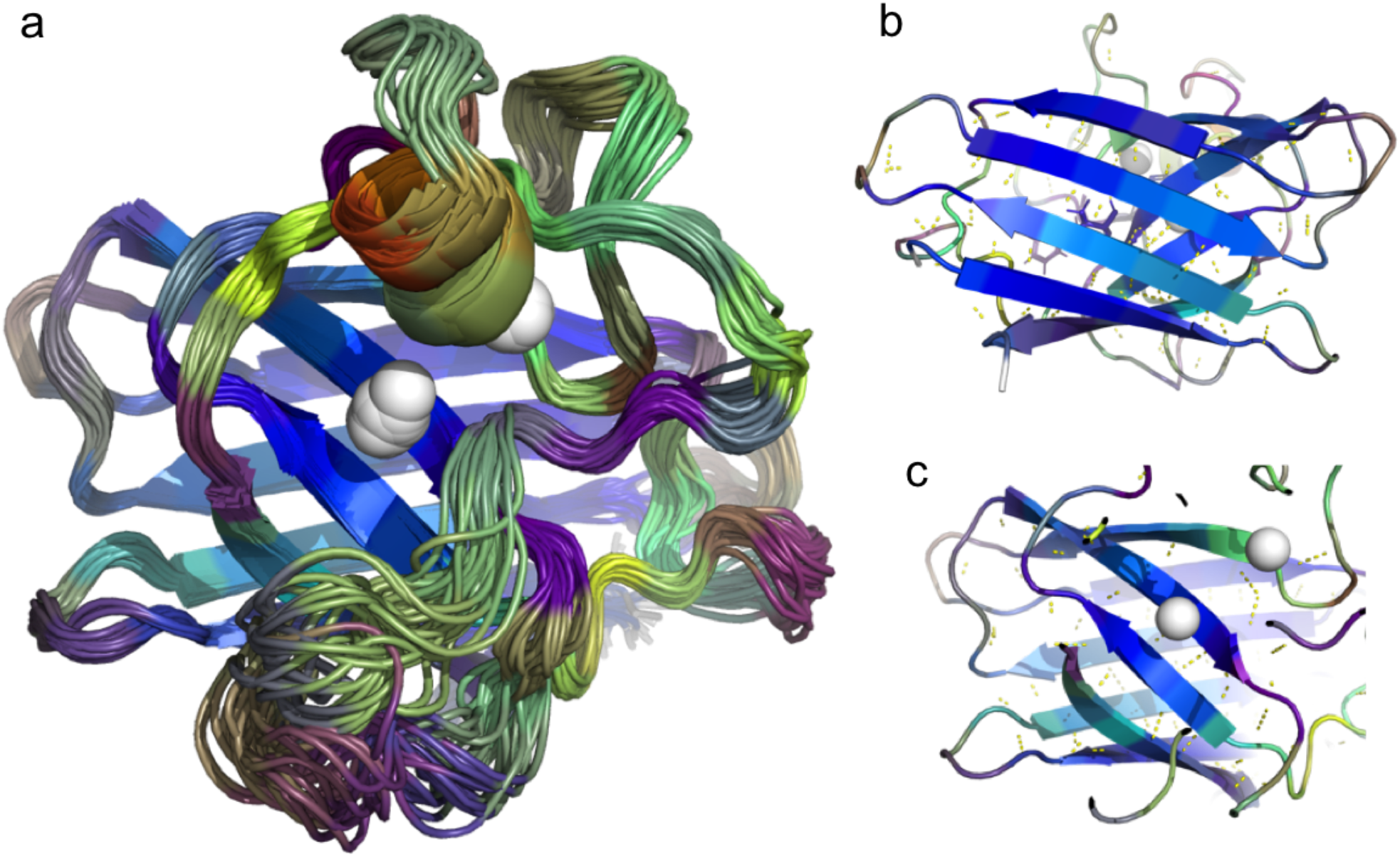
NMR structure ensemble of the Cu/Zn bound SOD1 colored with CheSPI colors derived from BMRB id 4402. **(a)** from “top” looking down the metal binding pocked. (**b**) from bottom facing strands 1, 2, 3, and 6. (**c**) from top “clipping through” the long loops to observe the top sheet of strands 4, 5, 7, and 8.

### Poly-proline II formation in an antifreeze protein

Poly-proline II helix (PPII) conformations (which are often, but not necessarily, rich in prolines) are relatively rare in folded proteins, although they appear to be important for certain molecular recognition events(Adzhubei, Sternberg et al. 2013). In contrast, PPII has been suggested to be more prevalent in IDPs(Shi, Chen et al. 2006) although it is not a generic conformation for IDPs, but part of the statistical composition of local structural states(Jha, Colubri et al. 2005, Makowska, Rodziewicz-Motowidło et al. 2006). δ2D predicts populations of the classical folded α-helix and β-sheet states, and populations of either so-called “coil” or “PPII”, which are both considered as disordered by δ2D. Similarly, CheSPI predicts “helical” and “extended” populations, but separates the remaining states into “turn” (folded DSSP Turn class) or “non-folded” (combined bend and coil DSSP states). In order to scrutinize the relationship between CSs and PPII, and in particular the CheSPI/CheZOD signatures possibly related to PPII, we analyzed spectroscopic and structural data for the left-handed helical bundle of Hypogastrura harveyi “snow flea” antifreeze protein (sfAFP), which is rich in Gly-Gly-X repeats(Graham and Davies 2005), and for which both an X-ray structure(Pentelute, Gates et al. 2008) as well as extensive NMR data including CSs are available(Pentelute, Gates et al. 2008). sfAFP has a compact structure of a bundle of six PPII helices connected by hydrogen bonds alternating between inter-strand neighbors with a three-residue periodicity (Fig. 9h). Relaxation measurement confirmed the rigid backbone - except for residues 25-31, which had elevated backbone dynamics. CheSPI analysis reveals some fluctuation in SCSs although with the absence of a common sign when scaled with the weight for the first component (Fig. 9a). This is reflected in Z-scores that typically lie between 10 and 12, which would indicate order, although the score is not as high as for fully formed standard helices and sheets (these lie around 13-15 typically). Residues 25-35 form an exception, and appear to be clearly more disordered, with Z-scores below 5. Furthermore, the PPII stretches feature CheSPI components much closer to zero (i.e. closer to random coil values with averages around ca. −2.0 and 0.0 for the first two components, and likewise near-zero secondary structure propensities by ncSPC) (Fig. 9b) than for ordered helices and sheets resulting in paler CheSPI colors closer to cyan-grey (Fig. 9c,i). The above ranges for CheZOD Z-scores and CheSPI components may be considered hallmarks of PPII helices, but with the current algorithm, CheSPI predicts primarily non-folded conformations for sfAFP and typically around 10-25% extended structure for the PPII stretches (Fig. 9d). In comparison, δ2D predicts around 25% PPII for these stretches (Fig. 9g), which is very similar to the PPII populations predicted for the IDPs tested here and for the case of human Tau protein discussed below (Fig. 10). The peptide segments forming PPII are apparent from the figure with exclusively coil DSSP classes (barring a few bends) and backbone angles in the PPII domain (Fig. 9 e,f). It could be suspected that the helical bundle in sfAFP might have peculiar and specific interactions, such as variations in twist along the PPII helices, as reflected here in the varying CheSPI colors, and both standard backbone and unusual Hα-C’ hydrogen bonds(Pentelute, Gates et al. 2008), that could potentially affect the CSs and thus the resulting CheSPI components. To address this systematically, our database of structured proteins was searched for consecutive stretches of three residues in PPII conformation (and other conformations, for reference) within DSSP “coil” states. Distributions of the CheSPI components for PPII in the database were found to be similar to the sfAFP case but were also found to be rather similar for stretches of three consecutive “extended” conformations (first CheSPI component was −0.97 and −1.64 in the former and latter case, respectively, i.e., with difference within one standard deviation, and the second component close to zero) (see Figure S4 and see also Figure 2). This was also the case for the PPII helix, residues 95-99, in the *P. aeruginosa* protein PA1324, protein (BMRB id 6343, see Figure S5). Hence, it remains very challenging to discriminate between “extended” (β-strand like) and PPII stretches using chemical shifts alone.

**Figure 9:**
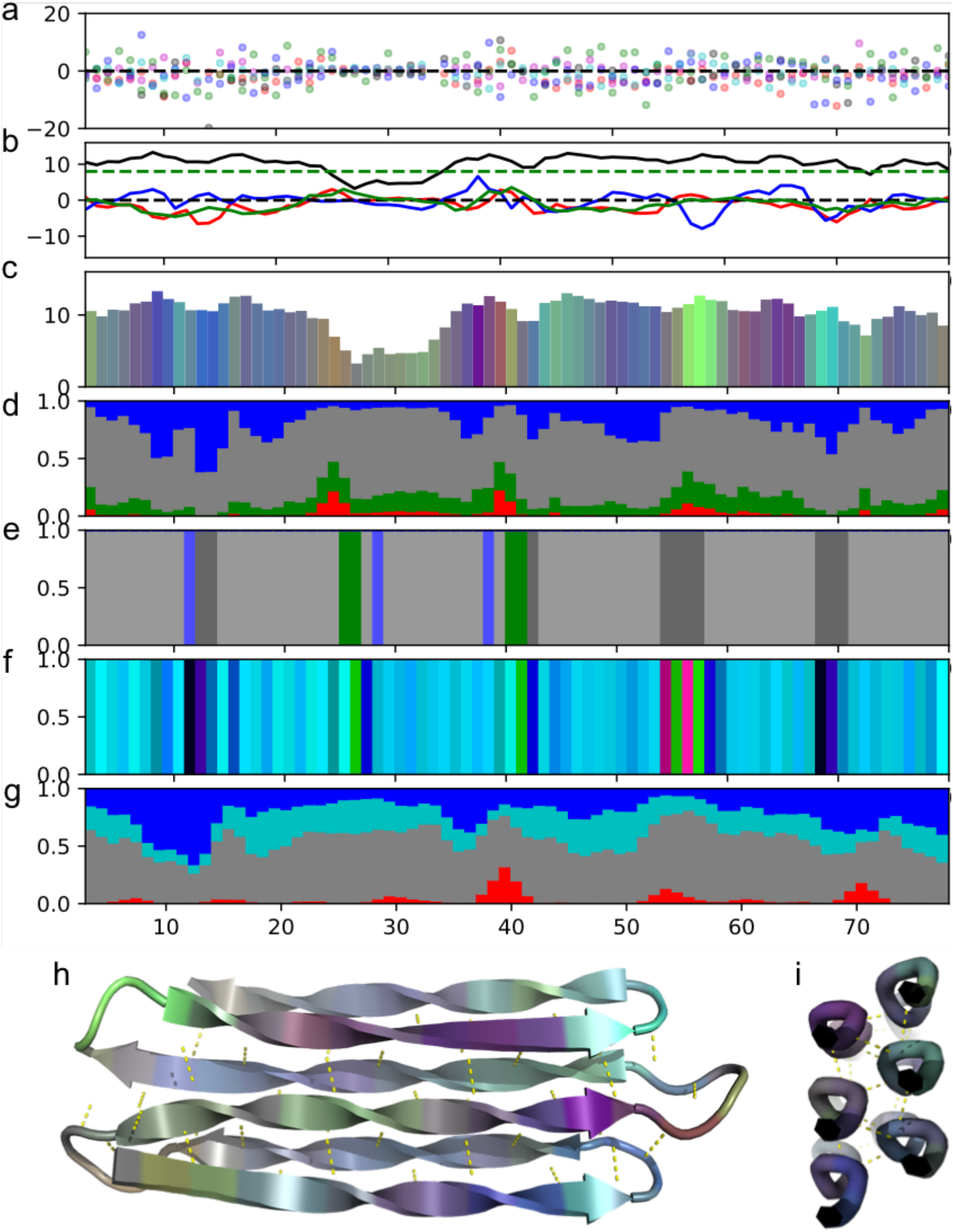
CheSPI analysis of PPII helical bundle protein, the Hypogastrura harveyi “snow flea” antifreeze protein (sfAFP) (**a-d**) CheSPI output derived from assigned chemical shifts for BMRB id 27473 (see legend to Figure S1 and Figure 4a,b) and the ncSPC secondary structure propensity multiplied with 8 is shown with a green broken line in panel b. (**f**) DSSP classification and (**g**) local backbone conformation in structure of sfAFP determined by X-ray crystallography with PDB id 2pne (see legend to Figure 4d,e). Predictions by δ2D visualized with stacked by plot showing helix, coil, PPII, and β-sheet using red, grey, cyan and red blue, respectively. (**h,i**) X-ray structure of sfAFP (PDB id 2pne) colored with CheSPI colors (as in panel c), hydrogen bonds are highlighted with yellow dashes. PPII and extended stretches are highlighted with standard β-strand cartoon rendering in (h) for visual purposes.

**Figure 10:**
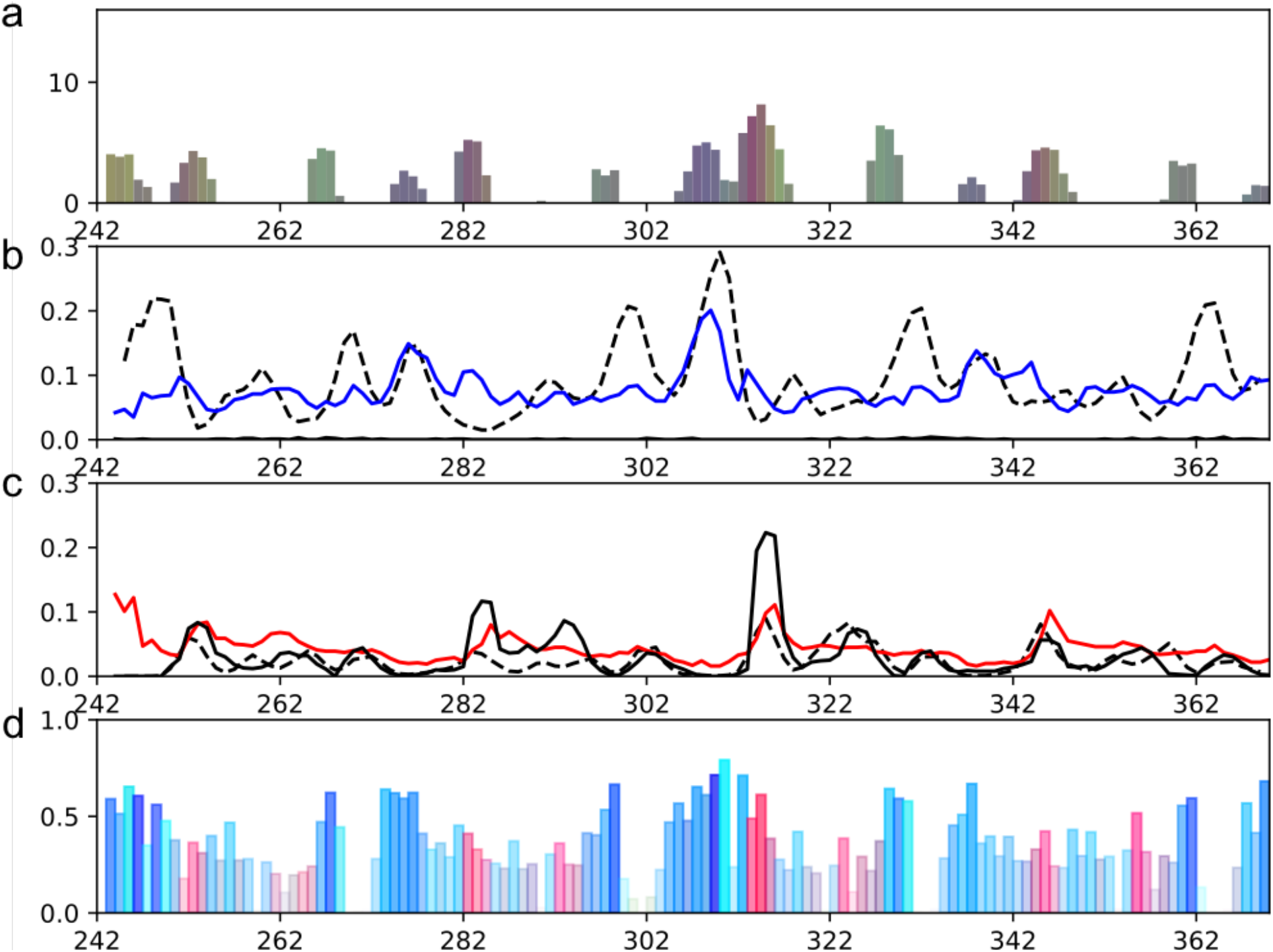
CheSPI output panels for K18-Tau and comparison to structural ensemble and δ2D. **a**) CheSPI colors based on assigned CSs from BMRB id 19253. **b**) CheSPI extended populations (blue curve), δ2D predictions of beta-strand (black broken curve), fractions of beta-strand or bridge formed in the ensemble structure of K18 Tau (black curve very close to zero) (PED id 6AAC). **c**) CheSPI helical populations (red curve), d2D predictions of helix (black broken curve), fractions of helix formed in the ensemble structure of K18 Tau (black curve). **d**) Average local backbone conformations in K18 Tau ensemble (see legend to Figure 4e). See also Figure S6.

### Aggregation nuclei of the protein Tau

IDPs are most fittingly interpreted as a statistical ensemble of local structural states. Here we demonstrate the ability of CheSPI to quantitatively infer the local structural composition from chemical shifts for a well-studied IDP, the K18 domain from human Tau (K18-Tau). Tau is intrinsically disordered, implicated in the regulation of microtubule organization, and prone to aggregation with a pathology related to Alheimer’s disease. Tau aggregates as neurofibrillar tangles, containing paired helical filaments (PHFs) that adopt cross-β and β-helix conformation akin to other amyloidogenic proteins(Berriman, Serpell et al. 2003, Barghorn, Davies et al. 2004). The K18 domain is directly involved in aggregation and consist of four imperfect 31-32 residue repeats, R1-R4. Hexapeptide segments within each repeat, residues, 275-280, 306-311, and 337-342, (HPF6, HPF6*, and HPF6**) are nucleation sites for aggregation, having a local β-sheet structure when forming multimers, and where HPF6 is at the core of the cross-β structure(von Bergen, Friedhoff et al. 2000, Eliezer, Barré et al. 2005, Daebel, Chinnathambi et al. 2012, Fitzpatrick, Falcon et al. 2017). K18-Tau was studied by NMR spectroscopy(Mukrasch, Markwick et al. 2007) and chemical shifts, RDCs, and PREs were used to calculate an ensemble of structures, accounting for the statistical composition of local structures, as outlined above(Ozenne, Schneider et al. 2012). An algorithm for efficient sampling of conformational space was applied, while simultaneously satisfying the diverse experimental constraints. Indeed, the simulations confirmed the unstructured nature of K18-Tau with very little regular secondary structure and revealed a mixture of compositions of primarily PPII and “extended” conformations, with fewer turns and helical formations. Although K18-Tau is largely disordered, the ensemble shows subtle sequence-specific variations in the local conformational composition with some similarities between its four pseudo-repeats.

Analysis by CheSPI (summarized in Figure 10 and Figure S6) reveals findings that agree very well with the earlier observations outlined above. SCSs display limited scatter (although with subtle variations) leading to low CheZOD scores < 8.0 indicative of disorder (Fig. S6a). Concomitantly, the first two CheSPI components show values close to zero, again with some variation. For comparison, the ncSPC-derived secondary structure propensity shows a close correspondence with the first CheSPI component (Fig. S6b). CheSPI predicts a preponderance of non-folded populations, albeit with important local biases (Fig. S6d). Firstly, elevated extended conformations are predicted by CheSPI for the hexapeptide HPF6(*/**) segments as shown in Figure 10b. It is interesting, that segments that are responsible for aggregation and part of the core cross-β structure are already more extended in the unfolded conformation. Secondly, shorter 3-4 residue segments following the FPF6s had higher CheSPI populations for helical structure (Fig. 10c). Indeed, turn structures were assigned to these segments measured from RDCs(Mukrasch, Markwick et al. 2007) DLKN (residues 253-256), DLSN, DLSK, and DKFD in repeats R1-R4). These turns are of type beta I, and two such consecutive turns correspond to a short 3_10_ helix(Pal, Chakrabarti et al. 2003). In the ASTEROIDS ensemble structure of K18-Tau (Fig. S6f), an excess of helical conformations and backbone angles were actually observed for these residues, in particular for the segment 313-315 in R3. Regular β-strand formation was not observed in the ensemble. However, a higher content of extended and PPII backbone conformations was encountered for residues in the HPF6 segments having also the highest CheSPI extended populations (Fig. 10d and Fig S6e). For comparison, δ2D (Fig. 10, black broken curve and Fig. S6g) identifies the same maximum for the helix conformation – but only with confidence for R3. Furthermore, δ2D identifies higher β-like conformation for the HPF6 segments, but also for other positions in the sequence that were found to have mixtures of helical, extended, and positive ϕ conformations (e.g. residues 298-303) in the ensemble derived from experimental data. The level of PPII conformation for K18-Tau predicted by δ2D (Fig. S6g) were also almost constant throughout the sequence and even slightly higher than the predictions for the PPII helical bundle discussed above.

### Misfolding of alpha-synuclein

The protein alpha-synuclein (aS) is disordered under native conditions, but prone to misfolding forming cytotoxic aggregates implicated in the pathogenesis of Parkinson’s disease(Singleton, Farrer et al. 2003, Stefanis 2012). One of the physiological functions of aS is its binding to synaptic vesicles where it adopts a semi-folded α-helical conformation(Davidson, Jonas et al. 1998, Jensen, Nielsen et al. 1998). This spurred a range of studies related to the binding of lipids and engineered membrane mimics(Jensen, Nielsen et al. 1998, Tofaris and Spillantini 2005). aS is comprised of seven 11-residue adjoined pseudo-repeats of amphiphilic character (I-VII) with a small flanking N-terminal four-residue insertion (between IV and V), and a longer acidic C-terminal region. A new variant with shuffled repeats, referred to as SaS, was designed previously to study the effect of sequence on vesicle binding and aggregation(Rao, Dua et al. 2008). aS and SaS was studied together with beta synuclein (bS) by NMR spectroscopy, analyzing the effect of interaction with sodium lauroyl sarcosinate (SLAS) micelles(Rao, Kim et al. 2009) and a structural model of aS-bound SLAS micelles was later derived based on NMR and EPR data(Rao, Jao et al. 2010). CheSPI analysis confirms aS to be disordered under native conditions (Fig 11a,e). In contrast, when interacting with SLAS micelles, all studied synuclein variants form structures with high helical content in the amphiphilic repeat region (Fig. 11b-d,f-h). Comparison with the ensemble structure model for aS reveals helix formation for residues 1-91 with partial interruption of the helical structure around repeat III and a small helix kink around residues 60-65 (Fig. S7c). The helix interruption region corresponds to the region with lowest CheSPI helical populations (repeat III, Fig. 11f) and the helix kink is located at a position in the sequence with less “canonical” CheSPI colors (end of repeat V, Fig. 11b), i.e. the colors transition to greener, which is found at the C-terminal end of a helix (see Results and Figure 4a) suggesting partial disruption of hydrogen bonding. Concomitantly, lower helical populations were found for the end of repeat V. A similar dip in CheSPI-derived helical populations and green colors were found at the end of repeat VII, although, in this case, no significant local irregularities were identified in the model structure (Fig. 11b,f). bS differs from aS by 14 mostly conservative mutations in the first 95 residues and the deletion of residues 73-83, disrupting repeats VI and VII. It is seen by the similarities of the CheSPI color profiles (Fig 11b,d) that bS, despite the sequence modification, retains the local structural and dynamical properties with e.g. similar transition to greener colors and deletion of residues 73-83 to display similar signatures for the end of the helical region. In analogy to the bS case, SaS and aS share the same positions for the last repeat VII, and similar CheSPI color profiles are observed, indicating near-identical local structural and dynamical properties. Furthermore, repeats I, II, IV and the insertion form canonical helices in aS as indicated by red CheSPI colors and close to 100% CheSPI helical population. Concurrently, the same repeats also show strong signatures of helix structure when repositioned in the SaS sequence (Fig. 11c,g). On the other hand, the full repeat III and the end of repeat V, which feature lowered CheSPI helical populations (about 50%), and have partially disrupted helical structure or a kink in the SLAS-bound model, also indicate lowered helical strength for SaS – but in this case about 75% CheSPI helical populations for repeat III and ca. 30% for the end of repeat V when repositioned in SaS. aS and its variants binds micelles due to the amphiphilic nature of their sequences (see Figure S7a,b). Repeats III and V contain more charged and hydrophilic residues and fewer amino acids with hydrophobic side chains, when compared to their neighbor counterparts, suggesting a lower micelle binding affinity (Fig. S7a,b), hence explaining the lower CheSPI helical populations. This exemplifies how the local structure in proteins and their interactions with substrates is mostly driven by the local sequence. The relocated repeats in SaS are modulated by the surrounding sequence/structure so that repeat III is presumably less helically interrupted whereas the end of repeat V is more kinked compared to aS.

**Figure 11:**
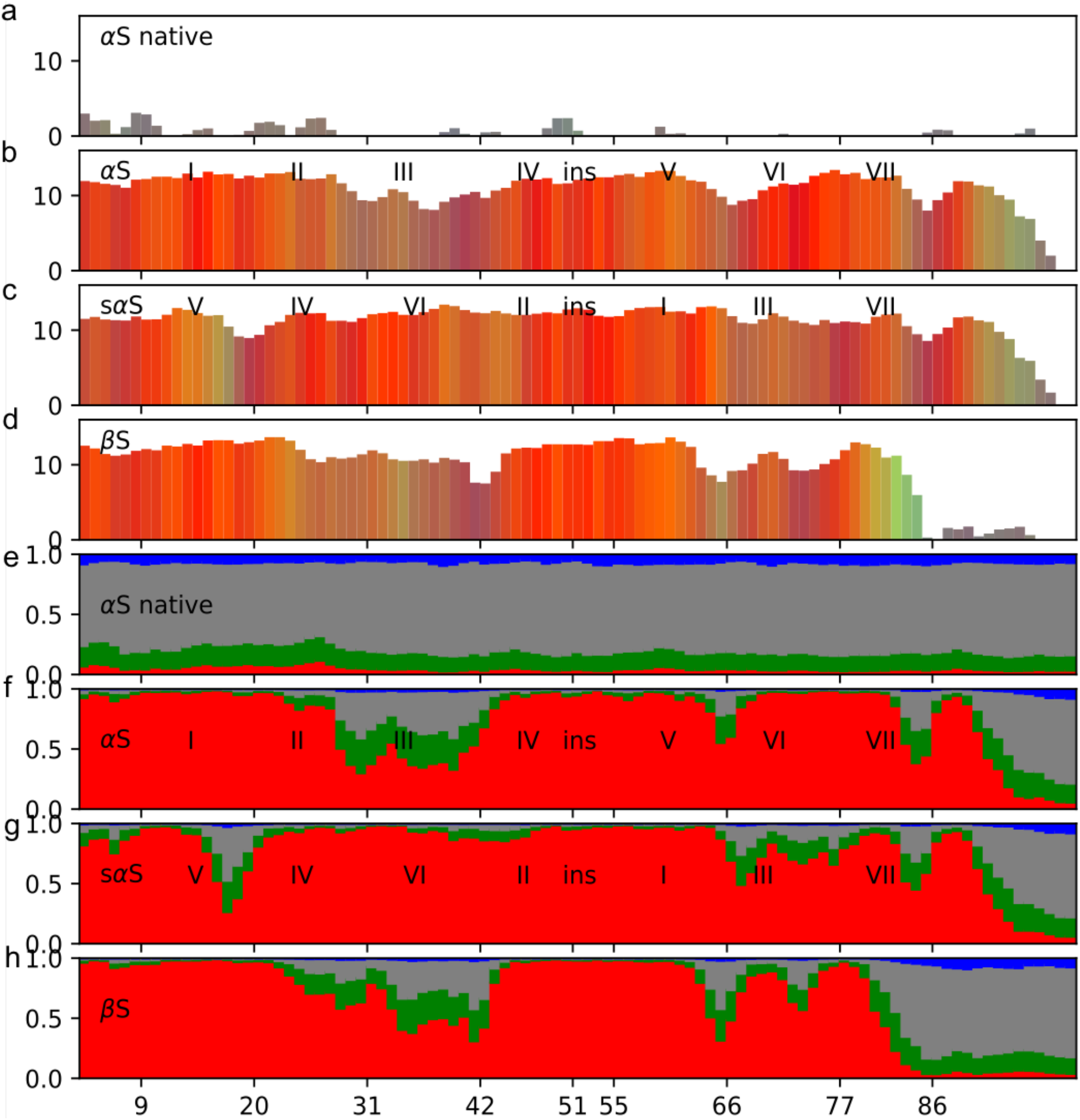
Alpha Synuclein variants. (**a**-**d**) CheSPI color bar plot (see legend to Figure 4c). (**e**-**h**) CheSPI populations stacked bar plot (see legend to Figure 4d). The individual panels show results for the following alpha synuclein variants and conditions: αSyn native disordered (BMRB id 16300 panels a and e), αSyn SLAS-bound (BMRB id 16302 panels b and f), “shuffled” αSyn (sαS, see text) SLAS-bound (BMRB id 16303 panels c and g) βSyn SLAS-bound (BMRB id 16304 panels d and h).

## Conclusions

We have introduced the software CheSPI for the comprehensive inference of structural and dynamical properties of proteins from assigned NMR chemical shifts and sequence. CheSPI can be applied to decompose chemical shifts to reduced dimensions and visualize protein secondary structure preferences using CheSPI colors. CheSPI provides predictions for the fine-grained conformations of local structure through estimated probabilities for the eight commonly recognized DSSP classes. A strong correlation was observed for Q8 (to recognize the eight classes solely from chemical shift data) and this was even stronger for Q3. It was demonstrated with a small number of examples how CheSPI can quantify and display the degree of protein disorder, and detect small populations of local structures in IDPs.

## Methods

### CheSPI components: the chemical shift principal components

The CheSPI component of order, *k*, for residue *i*, is computed as the weighted sum of truncated secondary chemical shifts (SCSs), Δ, for a 3-residue window as:

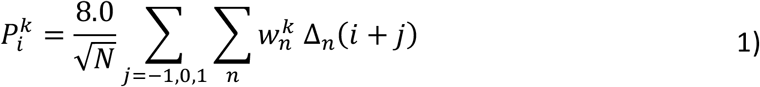

where

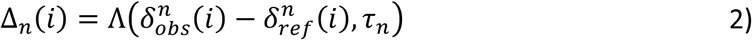

and

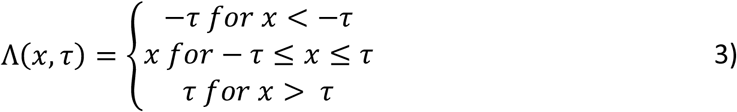

and τ_n_ are nuclei-specific values used to truncate SCSs to avoid extreme values caused by assignment errors or typos. Here *N* is the total number of available experimental CSs for the residue triplet and 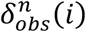 and 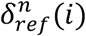 are the observed and reference CSs, respectively, for residue, *i*, and nuclei, *n*, where the reference CSs are the “random coil” CSs computed by POTENC(Nielsen and Mulder 2018). Note that the universal weight of 8.0 was chosen arbitrarily to obtain component values comparable to the CheZOD Z-score ranges. The weights, 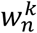, were derived by an Orthogonal Partial Least Squares Discriminant Analysis (OPLSDA) using SIMCA Umetrics(Wu, Li et al. 2010). This process identifies the linear combinations of the CSCs that best discriminates between the different secondary structure classes.

### CheSPI colors

The first two CheSPI components are visualized as a unique color, first scaling a component, x, to be between 0 and 1 and truncated between threshold values ±τ using

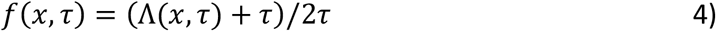

The color is then defined in terms of an RGB fraction vector, *C*, as:

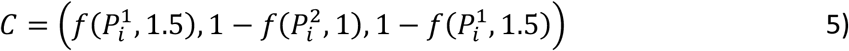

This definition leads to primarily blue colors for sheets, red for helices, and green colors for turns whereas disordered states with principal components close to zero correspond to grey colors.

### CheSPI populations: secondary structure populations inferred from CheSPI components

Helix, sheet and coil all have rather distinct distributions of CheSPI components as seen for the correlated distribution for the 809 proteins with positive and negative values for the first component for helices and sheets, respectively, whereas there is more overlap for coil states with average values for both components near zero. The population on of a 3-state secondary structure type, *s*, is calculated based on the density, ρ_s_, from the experimental correlated distribution of the two first CheSPI components:

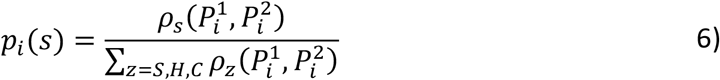

The densities were estimated from histogram distributions from the 809 proteins set.

### Prediction of 8-state DSSP secondary structure classes from sequence and CheSPI components

A back-calculation prediction model was defined for the CheSPI component, *P*, from local primary sequence and 8-state DSSP secondary structure class. The model is linear with a constant term corresponding to the DSSP class and corrections for the sequence, *C*, and secondary structure, *D*, values in a sliding window of four residues in each direction as:

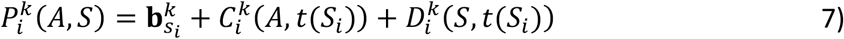

where

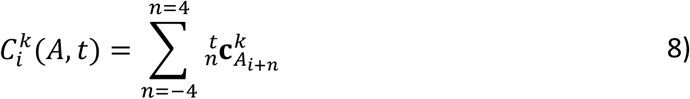

and

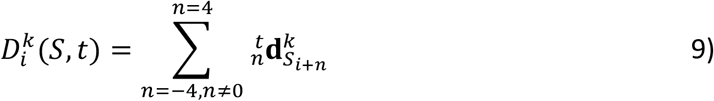

where A_i_ denotes the amino acid sequence, and S_i_ and T_i_ the 8- and 3-state secondary structure types, respectively, all at residue position *i,* and s and t denote position-specific 8- and 3-state secondary structure classes, respectively, and *t* maps the 8-state classes to 3 classes helix/sheet/coil. The correction term, C, constitutes 540 predetermined constants for each CheSPI component whereas the secondary term, D, constitutes 192 constants. These constants were derived from a multi-linear regression fit using the large set of 809 protein sequences with known secondary structure. Some constants were set to zero in order to limit the number of free parameters. The optimal balance between free parameters and goodness of fit was derived by minimizing Akaike’s Information Criterion (AIC) and varying the number of adjustable parameters using a genetic algorithm as described previously when parametrizing POTENCI(Nielsen and Mulder 2018).

The DSSP 8-state secondary structure classes are predicted using comparison between observed and back-calculated CheSPI components and by applying Bayes Theorem. First, based on the observed CheSPI components, *Q*, the likelihood, *L*, of observing the principal components, given a certain secondary structure and the sequence, is calculated as:

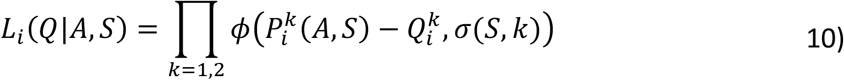

where ϕ(x, s) is the normal distribution density function with mean 0 evaluated at x with variance s^2^, and σ is the standard deviation of the prediction errors measured for the training set of the secondary structure S. Secondly, the posterior probability for the secondary structure is calculated by multiplying the above likelihood with a prior probability for the secondary structure:

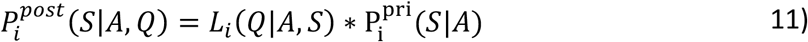

in other words, we use the CheSPI component data to update the prior probabilities for the secondary structure. The CheSPI application offers two procedures for estimating the prior probabilities: (*i*) simple per residue type frequencies for secondary structure types or (*ii*) secondary structure prediction based on sequences alone using Xraptor(Wang, Zhao et al. 2011, Källberg, Wang et al. 2012) Subsequently, the posterior probabilities for all 8 secondary structure classes are normalized to sum to unity.

Finally, the secondary structure predicted by CheSPI is identified as the configuration with maximum combined posterior probability for all residues:

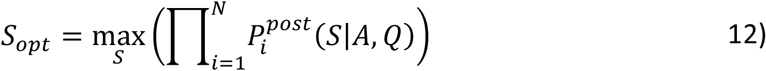

This problem has a large dimensionally and cannot be optimized exhaustively, and therefore, the secondary structure optimization was implemented with a genetic algorithm solver as was the case for POTENCI(Nielsen and Mulder 2018). The algorithm is initiated with random secondary structure conformations sampled based on the secondary structure predictions from sequence.

### Definition of backbone torsion angle averages order parameters

The dihedral angle order parameter, *S*, of Hyberts, Wagner and co-workers(Hyberts, Goldberg et al. 1992) is defined by averaged trigonometric values for backbone torsion angles:

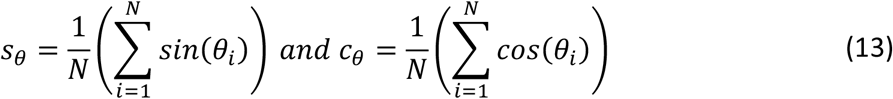

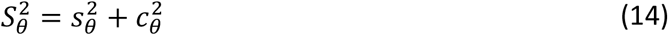

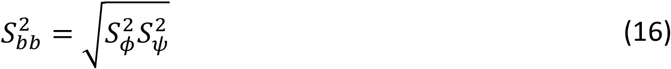

for an ensemble of *N* structures, where θ_i_ is the value of a particular dihedral angle θ in the *i*^th^ member of the ensemble and *Sbb* is the combined order parameter for the backbone torsion angles ϕ and ψ. The corresponding averaged angles are found by renormalizing with the order parameter and calculating inverse cosine:

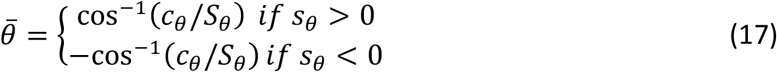

## Data availability

The datasets generated during and/or analysed during the current study are available from the corresponding author on reasonable request. Python code for CheSPI is available for download at GitHub: https://github.com/protein-nmr.

