## Supplementary Materials for "CheSPI: Chemical shift Secondary structure Population Inference"

### Content

Supplementary Results 1

Supplementary Figures S1-7

Supplementary Table 1

### Supplementary Results 1

CheSPI was applied to compute secondary chemical shift (SCSs( and CheSPI components using assigned chemical shifts for the phospholipase c epsilon RA 2 domain (Hyberts, Goldberg et al. 1992) (PLCε-RA2, henceforth). The analysis is summarized in Figure 4 in the main text and Figure S1 here. PLCε-RA2 contains 5 β-strands, two α-helices and two shorter $3_{10}$-helices, which are modelled in some of the members of the deposited NMR ensemble (PDB id 2byf). Figure S1a visualizes the SCSs for each residue. The fluctuations in SCSs for the structured parts of the protein are apparent, but are considerably smaller for the terminal residues and section corresponding to a very flexible loop in the structure. The trends described above for secondary structure are clearly visible, with high SCSs for Cα and C’ in helices and low SCSs for Hα and Cβ for β-sheets, in particular.

PLCε-RA2 feature high values of the first CheSPI component, $P^1$, for segments in helices, low for sheets and near-zero for both components for the flexible loop and around residue 70, and at the termini (Fig. S1b). The flexible loop also shows low CheZOD Z-scores, indicative of disorder. $P^1$ has somewhat lower amplitudes at the ends of the secondary structure elements. Some variation is observed in $P^2$ along the secondary structure elements, in particular starting with high values for the N-terminal ends of helices and ending with negative sign. The loop segment residues 43-56 has intermediate CheZOD Z-scores and lower amplitudes of $P^1$, indicative of dynamic or fractionally formed helices.

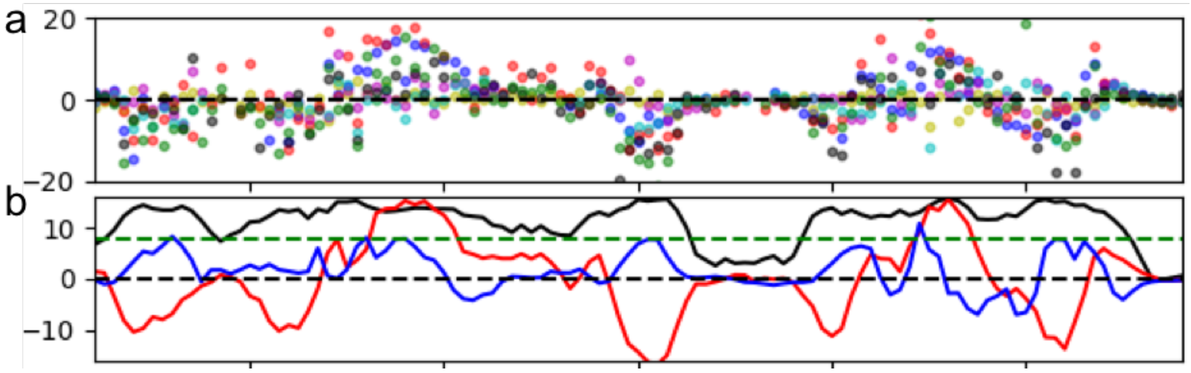


**Figure S1**: **Visualization of basic CheSPI and CheZOD output for PLCε-RA2 along the primary sequence** (**a**) Secondary chemical shifts (eq. 2 main text) multiplied with nuclei-specific weights (eq. 1 main text) to (for uniform variance) shown with blue, red, black, green, cyan, magenta and yellow dots for C´, Cα, Cβ, Hα, $H_N$, N and Hβ, respectively. (**b**) First two CheSPI components (Eq. 1 main text) using red and blue for the first and the second components, respectively, and the CheZOD Z-score shown as a black curve. The line of Z = 8 is shown for reference to indicate tentative border between primarily ordered and disordered residues.

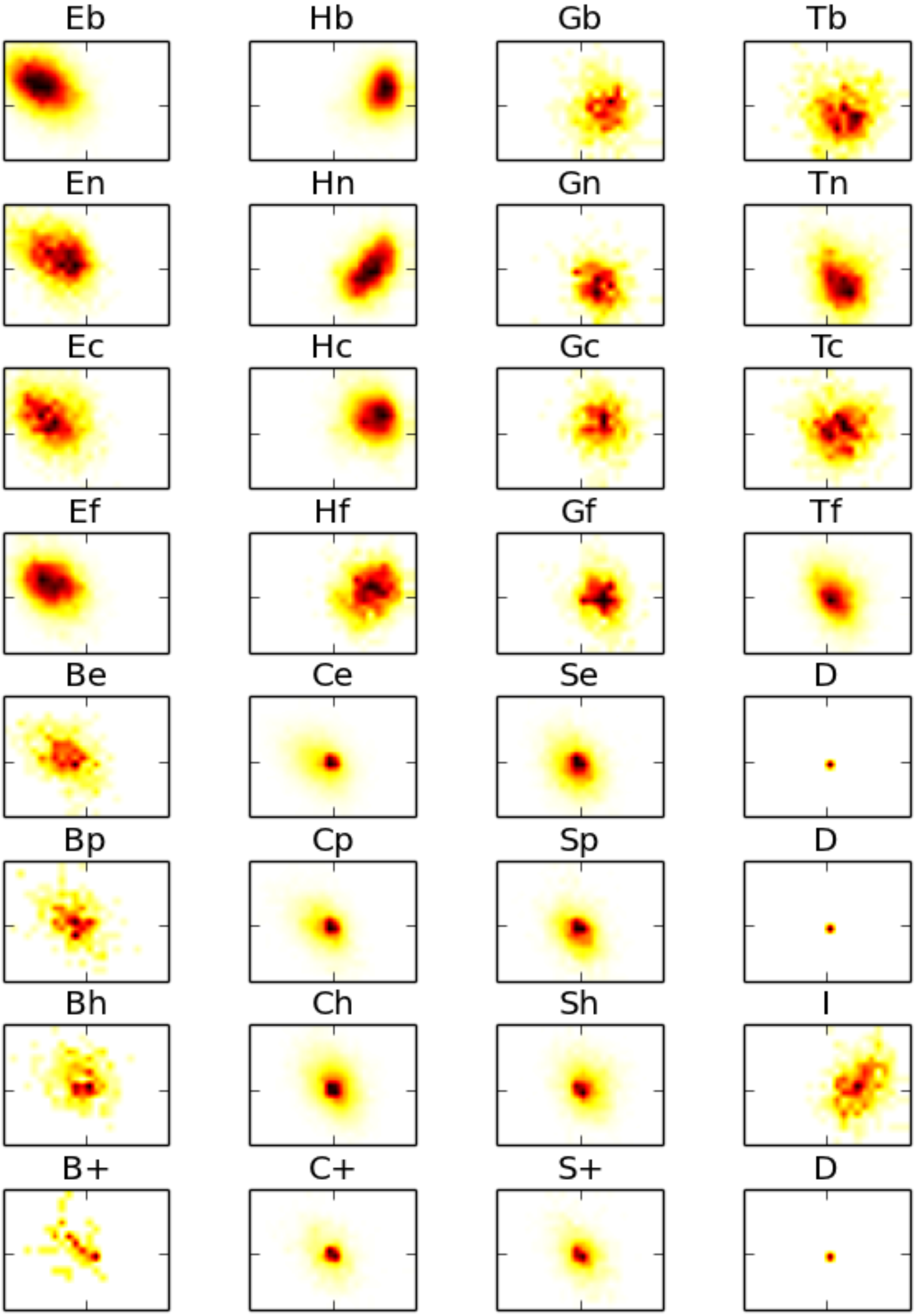


**Figure S2**: Histograms for correlated observations of the first two CheSPI components for all 8 DSSP secondary structure classes with subdivisions (see legend to Figure 3 main text and Table 1). Middle axis ticks mark $P^1 = 0$ and $P^2 = 0$, axis limits are -12/12 for the x-axis and -8/8 for the y-axis.

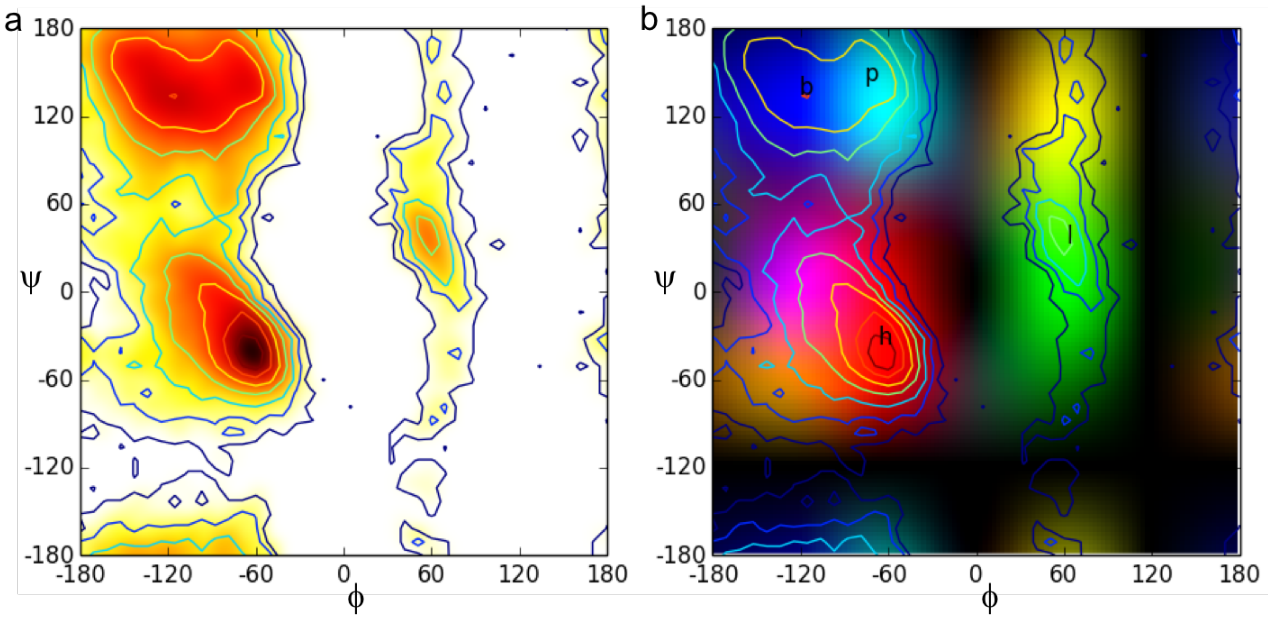


**Figure S3**: Ramachandran maps computed using the database of structured proteins showing ψ vs ϕ. (a) histogram showing higher densities with darker colors and contours superimposed. (**b**) 2D color-image with Ramachandran contours from (a) superimposed. Regions corresponding backbone angles in the helical domain ("h") of the Ramachandran map appear in red, extended β-sheet-like conformations ("b") have blue colors, left-twisted β-strands as well as fragments with PPII structure ("p") appears with cyan colors whereas conformations with positive ϕ ("l") have yellow and green colors, and finally, other conformation referred to elsewhere as "forbidden" in the Ramachandran space are have colors close to black. These colors applied elsewhere in some figures in the main text to visualize specific backbone conformation through their color correspondence (see e.g. Figure 4e in the main text).

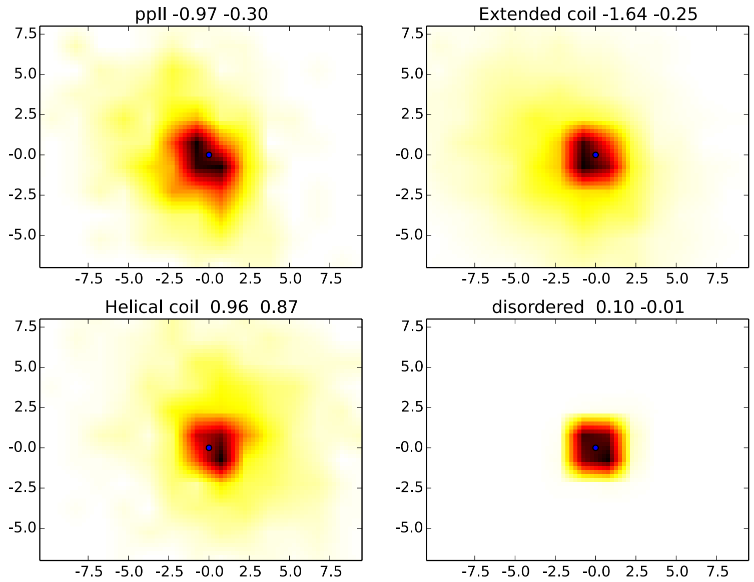


**Figure S4**: Histograms for the correlated distributions of the first two CheSPI components for PPII, extended, and helical coil for three consecutive residues in the structured database. Plot headers include the conformation followed by the mean values for the first two CheSPI components.

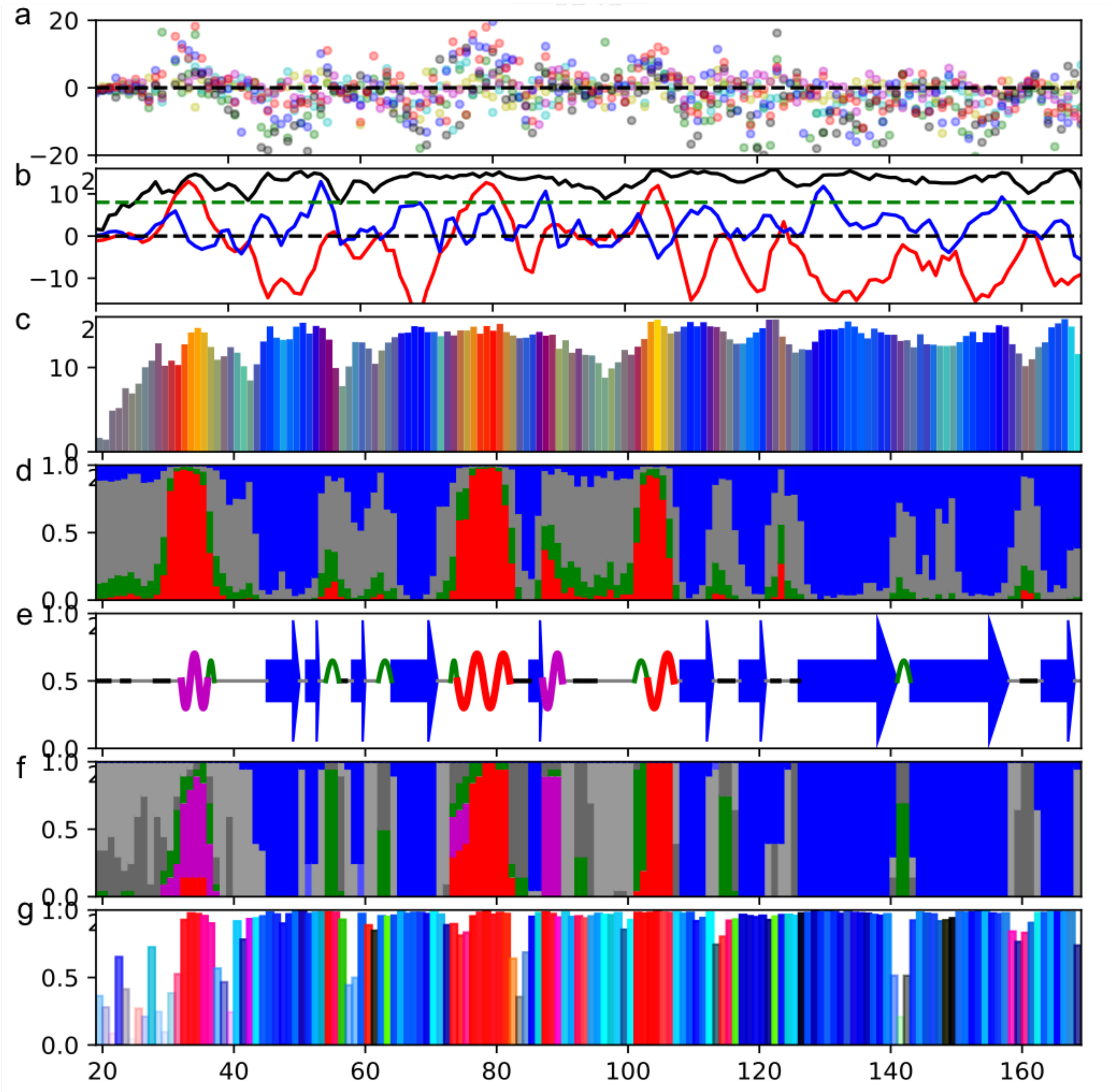


**Figure S5: CheSPI output** derived from assigned chemical shifts for BMRB id 6343, residues 21-170 (see legend to Figures 4 and S1) and ensemble local conformations for P. aeruginosa protein PA1324, residues 1-162, PDB id 1xpn (see legend to Figure 4d,e).

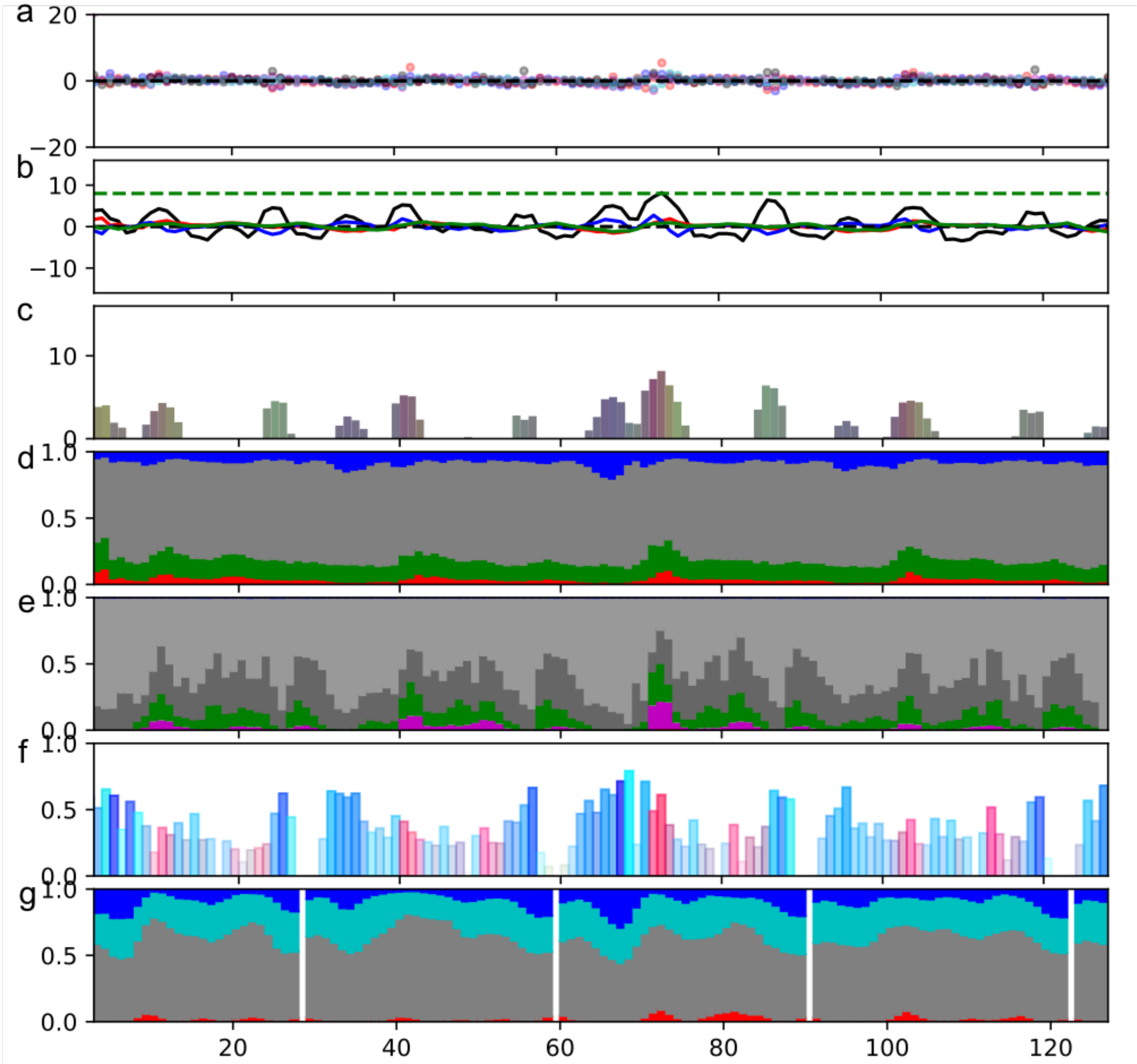


**Figure S6**: (**a**-**e**) **CheSPI full analysis for K18-Tau** (see legend to Figure 4a-e) **and comparison to structural ensemble, δ2D and ncSPC.** based on assigned chemical shifts from BMRB id 19253. The secondary structure propensity is shown with a green curve in panel b. (**e**) Stacked bar plot as in visualized in Figure 4d of ensemble conformation variation metrics, and (**f**) average backbone angle conformations as shown in Fig. 4e in the main text, using coordinates corresponding to p-ED id 6AAC (see main text) (**g**) δ2D predictions visualized as in Figure 9g.

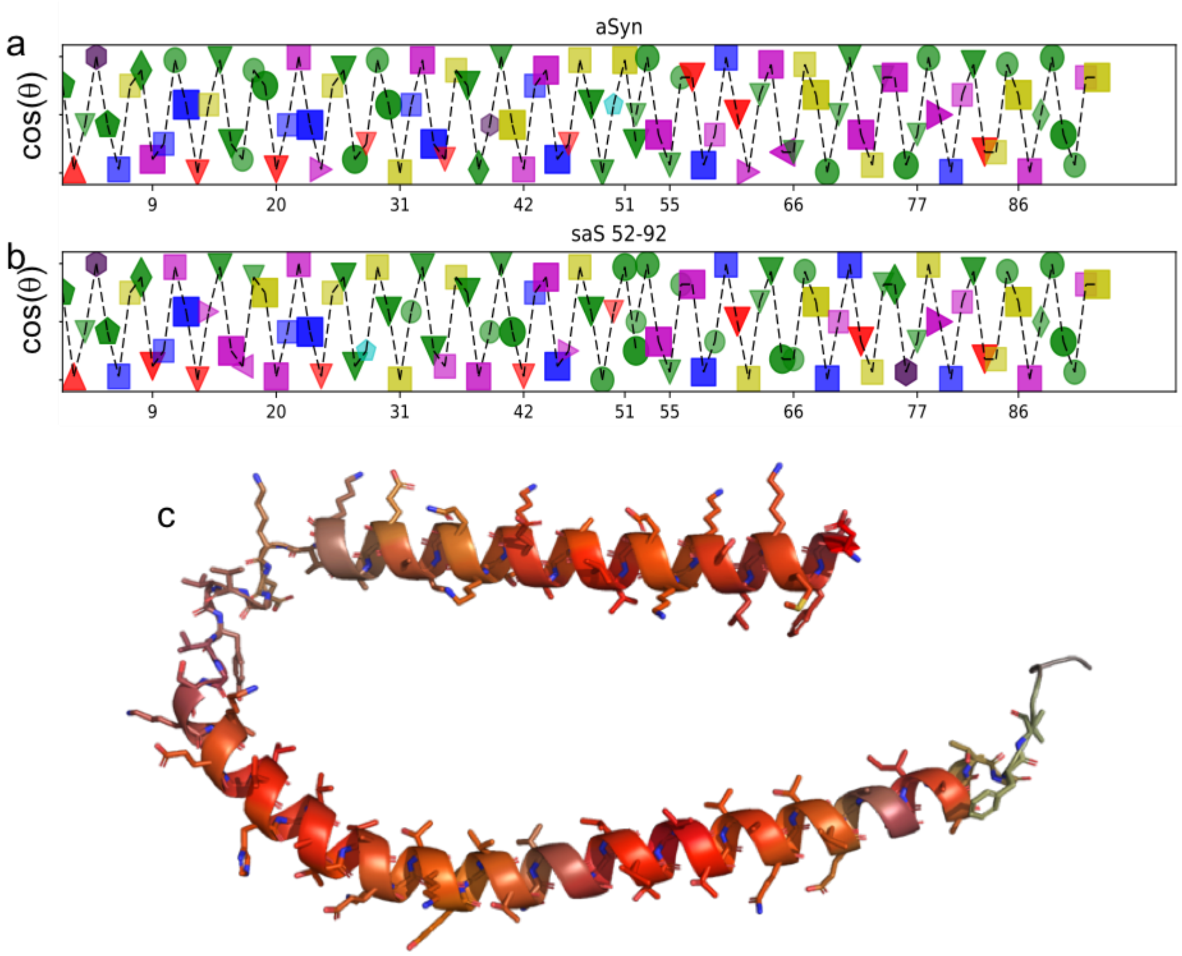


**Figure S7: Helical projections of alpha synuclein variants and cartoon model.** (**a**) alpha synuclein (aS) (**b**) shuffled aS (SaS) (see main text). (a,b) The helix is projected from the side with residue numbers shown on the x-axis. Sequence neighboring residues are connected with broken lines and each amino acid type is visualized with symbols, A(circles), STGK(squares), YF(hexagons), W(octagon), IL(diamonds), V(triangle pointing up), RHM(pentagons), and D/E/N/Q for triangles pointing up/down/left/right, respectively, with colored fill: G/P(yellow/grey), positive (blue), and negatively charged (red), aromatic(grey), His(cyan). (**c**) Structure model of alpha synuclein bound to SLAS micelles with residues colored using CheSPI colors (see Fig. 11b, main text).

**Table S1: Performance evaluation of CheSPI secondary structure predictions for 13 high quality protein structures**

| BMRB id | PDB id | $N_{res}$ | $f_{DIS}$ | Resolution/Å | R-value | Q3 CheSPI | Q8 CheSPI | Q3 CSI 3.0 | Q3 difference |
|---|---|---|---|---|---|---|---|---|---|
| 25379 | 2VU4A | 188 | 0.154 | 1.80 | 0.178 | 0.884 | 0.779 | 0.826 | 0.058 |
| 18574 | 1ISPA | 179 | 0.000 | 1.30 | 0.192 | 0.823 | 0.630 | 0.762 | 0.061 |
| 25354 | 1JO8A | 58 | 0.000 | 1.30 | 0.145 | 0.726 | 0.532 | 0.677 | 0.048 |
| 17586 | 3PO8A | 98 | 0.020 | 1.50 | 0.191 | 0.800 | 0.630 | 0.800 | 0.000 |
| 19282 | 3M9ZA | 137 | 0.080 | 1.70 | 0.197 | 0.791 | 0.612 | 0.806 | -0.014 |
| 18505 | 3HD4A | 136 | 0.037 | 1.75 | 0.189 | 0.783 | 0.594 | 0.826 | -0.044 |
| 25149 | 4R2YA | 68 | 0.015 | 1.75 | 0.195 | 0.886 | 0.586 | 0.871 | 0.014 |
| 18387 | 4GRFA | 150 | 0.040 | 1.76 | 0.168 | 0.901 | 0.770 | 0.836 | 0.066 |
| 17938 | 2HQHA | 86 | 0.140 | 1.80 | 0.194 | 0.841 | 0.648 | 0.886 | -0.046 |
| 18341 | 3ZDMD | 77 | 0.039 | 1.80 | 0.189 | 0.837 | 0.772 | 0.880 | -0.044 |
| 18029 | 3O55A | 119 | 0.000 | 1.90 | 0.189 | 0.915 | 0.767 | 0.884 | 0.031 |
| 18249 | 2B1KA | 173 | 0.093 | 1.90 | 0.190 | 0.909 | 0.806 | 0.909 | 0.000 |
| 19818 | 1Q1FA | 149 | 0.000 | 1.50 | 0.210 | 0.901 | 0.795 | 0.676 | 0.225 |
| | | | | | averages: | 0.846 | 0.686 | 0.818 | 0.027 |
